# Endophilin-Lamellipodin-VASP, key components in fast endophilin-mediated endocytosis, control actin polymerization within liquid-like condensates

**DOI:** 10.1101/2024.03.21.586200

**Authors:** Karthik B. Narayan, Honey Priya James, Jonathan Cope, Samsuzzoha Mondal, Laura Baeyens, Francesco Milano, Jason Zheng, Matthias Krause, Tobias Baumgart

## Abstract

Actin polymerization is essential in several clathrin-independent endocytic pathways including fast endophilin mediated endocytosis (FEME), however the actin machinery involved in FEME has been elusive. Here, we show that the actin polymerase VASP colocalizes and interacts directly with the FEME priming complex. We identify Endophilin as a VASP binding partner and establish novel non-canonical interactions between Endophilin and VASP. The major FEME regulators Endophilin and Lamellipodin interact multivalently with VASP to form liquid-like condensates both in solution and on lipid membranes that localize actin polymerization with the extent of actin polymerized regulated by multivalent Endophilin-Lamellipodin interactions. We identify a novel function for Endophilin condensates in bundling actin filaments and show that Endophilin directly binds filamentous actin. Our findings support a model that explains the connection between local actin polymerization and dynamic formation and dissolution of endocytic priming patches in FEME.

## Introduction

Endocytosis mediates the internalization of transmembrane proteins and of nutrients that are too large to diffuse through the plasma membrane. Numerous distinct endocytic pathways exist and can be categorized based on their dependence on clathrin, a membrane scaffolding protein, into clathrin mediated endocytosis (CME), and clathrin independent endocytosis (CIE). While CME is the prominent form of endocytosis in eukaryotes^1^, CIE routes function in parallel to internalize specific cargo^2^. CIE pathways like FEME are responsible for the uptake of activated receptors at the leading edge of cells to support migration^3^. In addition to being used for the exclusive internalization of certain receptors including β1-Adrenergic receptor (β1AR), and of the Epidermal Growth Factor Receptor (EGF receptor) when stimulated with elevated concentrations of EGF, it also acts as a pathway for rapid cargo uptake^3^. The successful internalization of cargo in both CME and CIE depends on the precise execution of multiple steps, which include the generation of membrane invaginations^1, 2, 4, 5^ and the local polymerization of actin^6–8^.

Actin polymerization is indispensable in most CIE pathways including FEME^3^, but the identity and mechanism of activity of the polymerization machinery and its interplay with endocytic regulators is poorly understood. Actin networks formed in-cellulo are a result of multiple polymerization machineries working in unison, often including nucleation promotion factors (NPFs) like N-WASP^9^, which activate the Arp2/3 complex and allow for the formation of branched actin networks, formins^10^, and Ena/VASP^11^ proteins that elongate actin filaments^12^. Interestingly, multiple studies have established a direct link between the membrane remodelling BAR (Bin, amphiphysin and Rvs) domain containing proteins and actin^13–15^ and its regulators^16–20^, prompting a need to understand the interplay between actin machineries^11, 21^ and membrane remodelling proteins^8, 16^ in membrane trafficking pathways.

In FEME, the BAR domain protein Endophilin (A isoforms) is recruited by Lamellipodin, an adaptor protein found at the leading edge, through interactions between Endophilin’s SH3 domain and the ten proline rich motifs (PRMs) found in Lamellipodin’s disordered C-terminal tail^3, 17^. Endophilin (referred to as Endo A1 or A2 depending on the experiment in figures and figure captions) exists asa homodimer formed via its BAR domain and contains an intrinsically disordered linker that connects the SH3 domain to the BAR domain (Fig 1A). Through multivalent interactions, Lamellipodin and Endophilin are enriched in transient protein patches (termed FEME priming patches) on the plasma membrane primed for endocytosis^3, 22, 23^. Along with its role in the formation of endocytic patches, Endophilin recruits dynamin, a downstream endocytic protein involved in membrane scission^24, 25^. Lamellipodin recruits Ena/VASP proteins to the leading edge of cells^26, 27^ and to clathrin-coated pits to support membrane scission^17^. We previously determined that Lamellipodin also recruits Endophilin A1-2^17^, and showed that Lamellipodin via Endophilin A1-2 and Ena/VASP proteins promote CME of the EGFR. Collectively, this prompted us to explore the possibility that Ena/VASP functions with Lamellipodin and both Endophilin A1 and A2 in FEME, in this study.

**Figure 1:**
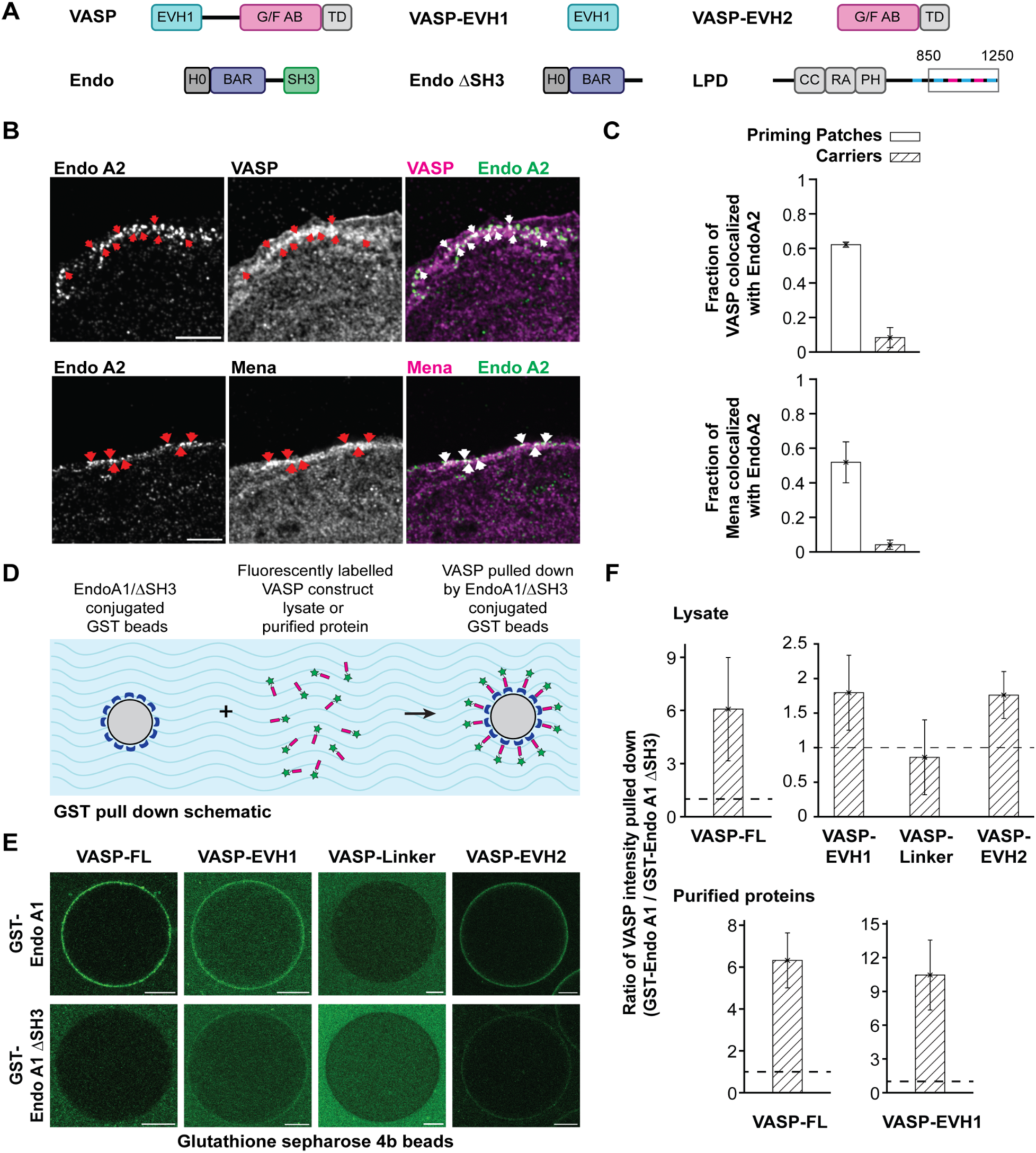
VASP colocalizes with FEME priming patches and directly interacts with Endophilin. **A** Schematic depicting the domains of Top row: VASP, EVH1 and EVH2. Bottom row: EndoA1, EndoA1 ΔSH3 and Lamellipodin (In box LPD^850-1250^). Blue and magenta strips in Lamellipodin’s C terminal tail represent the SH3 and EVH1 binding sites. **B** Endogenous colocalization of the Ena/VASP proteins, VASP and Mena with Endo-A2 at the leading edge of resting BSC1 cells. Scale bar 5 μm. **C** Fraction of FEME priming patches and carriers colocalizing with Top VASP, Bottom Mena. N= 15 cells from 3 independent experiments. The error bars correspond to the standard deviation. **D** Schematic of the experimental design of the pulldown of VASP and its domains by GST tagged EndoA1 or EndoA1 ΔSH3. **E** Pull down of VASP by EndoA1. Images show fluorescence signal of mScarlet-VASP (or its truncates) on Glutathione Sepharose 4B beads coupled with either 25 μM of GST-EndoA1 (Top row) or GST-EndoA1 ΔSH3 (Bottom row). Lysate prepared from HeLa cells expressing mScarlet VASP (or truncates) was incubated with EndoA1 or EndoA1 ΔSH3 beads for 30 minutes. Beads were washed and imaged. Scale bar 20 µm. **F Top** Quantification of fluorescence intensity shown in D. Bar graph shows the ratio of the fluorescence intensity of mScarlet-VASP (or its truncates) on EndoA1 coupled Glutathione Sepharose 4B beads to EndoA1 ΔSH3 coupled beads. Dashed line represents ratio = 1. N > 10 for all experiments. **Bottom** Quantification of pull-down of purified VASP or VASP-EVH1 by EndoA1. 5 µM VASP or VASP-EVH1 with 250 nM Alexa 594 labeled protein was incubated with GST-EndoA1 or GST-EndoA1 ΔSH3 coupled Glutathione Sepharose 4B beads for 30 minutes. Beads were washed and imaged. Dashed line represents ratio = 1. N > 10 beads for all experiments.

The Ena/VASP protein family, Mena, VASP, EVL in vertebrates, are homotetramers with each monomer comprised of an EVH1 and EVH2 (Ena/VASP Homology 1 or 2) domain connected by a disordered linker (Fig 1A). This proline rich linker harbours profilin-actin binding sites^28^, and enables interactions with SH3 domain containing proteins like c-Abl^29^ and IRSp53^18^. Ena/VASP proteins are recruited to the leading edge by the binding of their EVH1 domains to the seven Ena/VASP binding motifs found in Lamellipodin’s C-terminal tail (Fig 1A). The EVH2 domain engages with globular and filamentous actin promoting a speed up of actin filament elongation by temporarily preventing capping of actin filament barbed ends through capping protein and by the recruitment of polymerisation competent profilin-actin monomers^11^. The enrichment of the Ena/VASP proteins at the leading edge of cells is often accompanied by formation of filopodia^18, 30–32^ or development of lamellipodial fronts^26^, conferring an important role for Ena/VASP proteins in cell migration. Amongst the Ena/VASP proteins, VASP has been studied in most detail.

Multivalency is a key feature of the interactions between VASP and its binding partners including Lamellipodin. Biomolecules that interact multivalently can result in the formation of liquid-like condensates by a phenomenon called liquid-liquid phase separation (LLPS)^23, 33, 34^. LLPS has been demonstrated to play a critical role in the organization and function of proteins in-cellulo as the liquid-like condensates offer a platform to locally enrich proteins by specific sequestration, and to regulate reaction kinetics^35–37^. Interestingly several actin and microtubule regulators^34, 38, 39^ including VASP^40^ undergo LLPS that can drive the rapid polymerization of cytoskeletal proteins. Furthermore, recent contributions by us and others have shown that endocytic proteins^23, 41–45^ can form liquid like condensates at various stages of endocytosis. Our understanding of the function of LLPS in endocytosis is developing, and likely is important for the generation of membrane curvature and the regulation of actin polymerization.

In FEME, the local polymerization of actin is essential for the formation of endocytic priming patches^3^. However, the identity of the actin polymerases involved and the role of actin in the formation of the priming patches is poorly resolved. In this study, we first examine the colocalization of VASP and Endophilin at FEME priming patches in-cellulo and ask if direct interactions between the two proteins exist. Using recombinantly purified proteins, we investigate the capacity of VASP to form liquid-like condensates with Endophilin and Lamellipodin, in the bulk and on lipid membranes, in-vitro. We then probe the ability of the condensates to localize actin polymerization and look to identify molecular interactions that regulate the degree of actin polymerized. We finally propose a model that explains the role of actin in the formation of priming patches in FEME.

## Results

### Ena/VASP proteins colocalized with FEME priming sites

Given lamellipodin’s key role in FEME^3, 22^, and its seven binding sites for Ena/VASP proteins, we first asked if Ena/VASP proteins colocalize with FEME priming patches at the leading edge of the cells. We probed the colocalization of VASP and Mena (both members of the Ena/VASP family) with the most ubiquitous isoform of Endophilin (A2) by co-immunostaining in BSC1 cells, which have been documented to exhibit robust FEME^46^. Indeed, both Ena/VASP proteins showed strong colocalization with Endophilin-A2 at FEME priming patches on the plasma membrane but not on endocytic carriers distal from the membrane (**Fig 1B, 1C**). This result suggests a possible role for Ena/VASP proteins in the early stages of FEME, namely the generation of priming patches and/or in membrane invagination and scission. Given the well established in-vitro characterization of VASP^12, 40, 47–50^ as compared to Mena, the in-vitro experiments described below were performed with wild type VASP and it’s truncates.

### Endophilin directly binds VASP

Lamellipodin facilitates the recruitment of VASP to the leading edge^12^. Additionally, through its proline rich linker, VASP interacts with SH3 domain containing proteins like IRSp53 and c-Abl^18, 29, 51^. To uncover additional possible mechanisms of the recruitment of VASP to the FEME priming patches, we tested the ability of Endophilin to interact with VASP (going forward Endophilin A1 is used in all in-vitro studies and will be referred to as Endophilin in the text and captions, and EndoA1 in the figure legends).

The following VASP constructs: VASP-FL, VASP-EVH1, VASP-EVH2, and the linker domain (**Fig 1A**), were expressed as fusions with a fluorescent mScarlet tag in Hela cells. We performed a pulldown using the lysates obtained from the cells expressing the various VASP constructs with Endophilin-FL as the bait and Endophilin-ΔSH3 as the control (Fig **1D**). The fluorescence intensities on the GST beads (**Fig 1E, Supplementary** Figure 1) were quantified, and a ratio of the mScarlet intensity on the beads coupled with Endophilin-FL to Endophilin-ΔSH3 was calculated (**Fig 1F, Supplementary** Figure 1 **for more details on the calculation**). Using this ratio, we were able to conclude that VASP and Endophilin interact with each other through Endophilin’s SH3 domain, and VASP’s EVH1 and EVH2 domains (**Fig 1F**). Unlike previous reports that had reported interactions of the linker with SH3 domain containing bacterially purified proteins (excluding Endophilin)^18, 29, 51, 52^, we did not observe significant interaction between the linker domain of VASP with the SH3 domain of Endophilin (**Fig 1D, 1E**). This could be explained by Endophilin’s SH3 domain not being a binding partner of the proline rich region within the linker. Alternatively, binding of SH3 domains to VASP’s linker obtained from mammalian cell lysate might be abrogated in our case by the phosphorylation of S157^53, 54^.

The cell lysates used in the pull-down contains proteins (including Lamellipodin) that could facilitate an indirect interaction between Endophilin and VASP; therefore, to confirm direct binding of VASP to Endophilin, we performed the above-described pull-down experiment using recombinantly expressed and purified (from E. coli) VASP, as well as VASP-EVH1 (Lamellipodin binding domain). Consistent with the cell lysate experiments, purified full-length VASP depicted SH3 dependent binding to Endophilin (**Fig 1F**). Interestingly, VASP’s EVH1 domain also showed direct binding to Endophilin’s SH3 domain. These results together help establish a novel binding interaction between VASP and Endophilin, and non-canonical associations between the EVH1 and EVH2 domains of VASP with the SH3 domain of Endophilin.

### Key FEME regulators undergo LLPS

Recent work from our group identified the capability of the key FEME protein Endophilin to phase separate by itself and with Lamellipodin^23^. Similarly VASP has also been shown to form condensates by itself ^40^. From our findings above, we have established VASP’s colocalization with FEME priming patches and direct binding to Endophilin. Could VASP’s enrichment at the priming sites be a result of its capacity to form liquid-like condensates with Endophilin and Lamellipodin?

Utilizing protein purified from *E.coli*, we investigated the phase separation behavior of Endophilin (FL), VASP (FL) and Lamellipodin (aa 850-1250, referred to as Lamellipodin^850–1250^ throughout the text and LPD^850-1250^ in figures), in vitro. The Lamellipodin^850-1250^ fragment has been previously used by us and others to study the protein’s function in-vitro ^12, 23, 32^. We generated binary mixtures of VASP with either Endophilin or Lamellipodin^850–1250^ in the bulk and observed the formation of liquid like condensates (**Fig 2A, 2D**). We examined the spatial distribution of the proteins in the condensates by including a minority fraction of fluorescently labeled proteins (250 nM Alexa 488 labeled Endophilin, Alexa 594 labeled VASP and Alexa 647 labeled Lamellipodin^850-1250^) along with unlabeled protein. We quantified the protein intensities in the interior and exterior of the droplets and found that Endophilin and VASP were ∼45x and ∼15x enriched respectively, and Lamellipodin^850-1250^ and VASP were ∼40x and ∼30x enriched respectively, within the protein condensates (**Fig 2B, 2E**).

**Figure 2:**
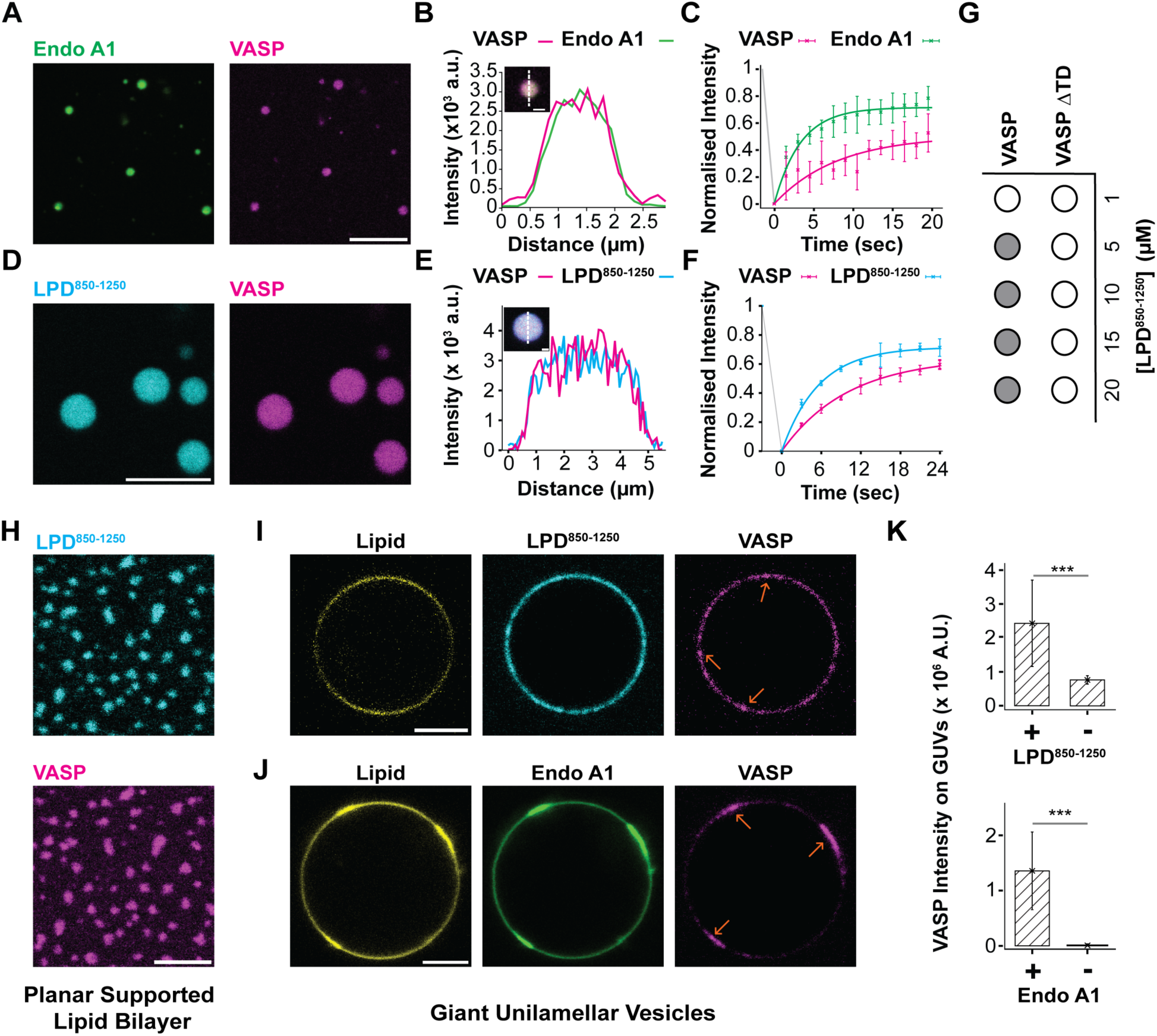
VASP forms liquid-like condensates with FEME priming proteins in the bulk and on lipid membranes. **A** Phase separated droplets formed by 30 μM each EndoA1 and VASP with 250 nM Alexa 488 and 594 labeled proteins respectively. Scale bar 5 μm. **B** Line intensity profile along red dashed line shown in inset image. Droplet generated under conditions described in **A**. Scale bar 1 µm. **C** Normalized FRAP profile of VASP and EndoA1 in condensates described in **A**. The error bars represent the standard deviation of 3 independent samples. Solid lines represent single exponential fits of the average recovery. EndoA1 Mobile fraction 0.71, Halftime of recovery 3.31 s, VASP Mobile fraction 0.50, Halftime of recovery 8.01 s. **D** Liquid-like condensates formed by 20 μM VASP and 10 μM LPD^850-1250^ doped with 250 nM Alexa-594 and Alexa-647 labeled VASP and LPD^850-1250^ respectively. (scale bar 5 μm). **E** Line intensity profile along dashed line drawn across LPD^850-1250^ -VASP (10 μM LPD^850-1250^ and 20 μM VASP with 10% labeled protein) droplet (inset). **F** Representative FRAP recovery curve for LPD^850-1250^ and VASP in condensates corresponding to conditions described in Left. The error bars correspond to the standard deviation of 3 independent measurements. Solid lines represent single exponential fits of the average recovery. LPD^850-1250^ Mobile fraction 0.71, Halftime of recovery 5.56 s, VASP Mobile fraction 0.64, Halftime of recovery 10 s. **G** Phase separation behavior of VASP and VASP-ΔTD with 10 μM LPD^850-1250^. Filled and open circles represent cases where droplets were and were not observed, respectively. At least three independent experiments were performed. **H** SLB (1% Ni-NTA DOGS, 99% DOPC) coupled with 100 nM His tagged LPD^850-1250^ after incubation with 2.5 μM VASP (doped with 250 nM Alexa 594 labeled VASP). Image acquired after 30 minutes of addition of VASP. Scale bar 5 μm. Independent repeats > 3. **I, J** Recruitment of VASP (1μM VASP with 10% Alexa 594 labeled VASP) to GUVs (45:30:24:1 mol% DOPS:DOPE:DOPC:MB-DHPE) in the presence of J LPD^850-1250^ (1 μM LPD^850-1250^ with 10% Alexa 647 labeled LPD^850-1250^) (**I**) or EndoA1 (2 μM EndoA1 with 5% Alexa 488 labeled EndoA1) (**J**). Scale bar 5 μm. Independent repeats > 3 **K** Intensity of VASP in clusters on GUVs corresponding to conditions described in **I, J** respectively. N>=10 GUVs per condition. Error bar corresponds to +/-1 standard deviation. Analysis: unpaired student t test was performed with a P value of < 0.005 (***).

Proteins in liquid-like condensates are expected to diffuse within the droplet. We quantified their fluorescence recovery after photobleaching (FRAP), to assess the translational mobility of VASP, Endophilin and Lamellipodin^850-1250^. We estimated the mobile fraction and characteristic diffusion times for the proteins in the condensates by fitting the recovery measurements to a single exponential function (**Fig 2C, 2F**). Specifically, VASP had a value of t_1/2_ ∼ 10 s for both condensate systems, and Endophilin and Lamellipodin^850-1250^ showed quicker recoveries with t_1/2_ of 3.3 s and 5.5 s respectively. The proteins in VASP-Endophilin condensates showed mobilities of 50% for VASP and 71% for Endophilin, and 64%-VASP and 71%-Lamellipodin^850-1250^ in VASP-Lamellipodin^850-1250^ droplets. These results together show that VASP forms liquid-like condensates in the bulk with Endophilin and Lamellipodin^850-1250^.

Recent reports have shown the essential role of VASP’s tetramerization domain in VASP’s capacity to form liquid-like condensates^40^. To assess the contribution of multivalency in the formation of protein condensates we generated a phase diagram of Lamellipodin^850-1250^ with VASP and the monomeric VASP construct (lacking the tetramerization domain), VASP-ΔTD. We formed binary protein mixtures of Lamellipodin^850-1250^ (10 µM) with both the full-length protein and VASP-ΔTD over a concentration range from 0-20 µM and confirmed that VASP-ΔTD was unable to form condensates with Lamellipodin^850-1250^ at all the conditions examined (confirmed by light microscopy) (**Fig 2G**). It has previously been shown that the phase separation behavior of the VASP-PEG system is controlled by electrostatics^40^. We therefore asked if electrostatics played a critical role in the formation Lamellipodin^850-1250^-VASP (without PEG, as used in Ref ^40^) condensates. To answer this question, we investigated the phase separation behavior of VASP and Lamellipodin^850-1250^ at increased ionic strengths. We observed that the capacity of VASP (20 µM) and Lamellipodin^850-1250^ (10 µM) to form condensates was impaired at 500 mM NaCl and was lost entirely at 1000 mM NaCl (**Supplementary** Figure 2). This indicates that Lamellipodin^850-1250^-VASP phase separation is dependent on electrostatic attraction between the molecules.

In FEME, endocytic priming patches are formed on the plasma membrane prior to receptor activation^3^. Our previous work^23^ has shown that it is possible to recreate endocytic protein rich clusters with in-vitro reconstitution models, using a supported lipid bilayer (SLB) system. We introduced VASP (2.5 µM total, 10% Alexa 594 VASP) to Lamellipodin^850-1250^ (100 nM, Alexa 647 labeled) coupled SLBs (99% DOPC/1% Ni-NTA DOGS) and observed the formation of protein clusters enriched in both VASP and Lamellipodin^850-1250^ (**Fig 2H, Supplementary** Figure 3A) mimicking FEME-like endocytic clusters. We next asked if multivalent interactions are essential for the development of clusters. To answer this, we replaced VASP with VASP-ΔTD and incubated it with Lamellipodin^850-1250^ coupled SLBs (same concentrations as above). In this condition, we did not observe the formation of spatially distinct VASP-Lamellipodin^850-1250^ clusters (**Supplementary** Figure 3B**, 3C**) providing further evidence for the indispensable role of multivalent interactions between the molecules in undergoing LLPS.

### Endophilin promotes lipid and protein clustering

Our finding that VASP rich clusters can be observed on neutrally charged membranes prompted us to investigate whether similar protein clusters can be formed on lipid bilayers that better mimic the inner leaflet of the plasma membrane. We asked if Lamellipodin^850-1250^ and Endophilin could recruit VASP to form clusters on lipid bilayers composed of 45:30:25 DOPS/DOPE/DOPC. Due to challenges incorporating conically shaped lipids (with inherent negative curvature), like DOPE^55^, into SLBs, Giant Unilamellar Vesicles (GUV) were utilized. We have previously shown that Endophilin can be recruited to bilayers of this composition^23, 56^ and we wondered if Lamellipodin^850-1250^ as well, due to its net charge of +17, could be recruited to this bilayer, by electrostatic attraction. We introduced Lamellipodin^850-1250^ to GUVs with and without DOPS (negatively charged) lipids and found that Lamellipodin^850-1250^ colocalized only with GUVs containing negative charge (**Supplementary** Figure 4).

We next introduced VASP (1μM total, 10% Alexa 594 VASP) to GUVs preincubated with either Endophilin (2 µM total, 5% Alexa 488 labeled) or Lamellipodin^850-1250^ (1 μM total, 10% Alexa 647 labeled) and observed VASP’s recruitment by both Endophilin and Lamellipodin^850-1250^ coupled GUVs (**Fig 2I-K**). Moreover, we noticed the formation of protein rich puncta on the membrane for VASP and Lamellipodin^850-1250^ (**Fig 2I**) and large protein domains on the GUVs incubated with VASP and Endophilin (**Fig 2J**). Interestingly, in the presence of Endophilin and VASP, we observed co-clustering of the lipid membrane with the protein domains (**Fig 2J**). The increase in the fluorescence signal could be explained by the formation of sub-resolution tubules resulting in apparent higher density of lipids. Such tubules have been shown to originate from phase separated protein domains on lipid bilayers^57^, ^58^. Alternatively, the increase in fluorescence intensity could be due to changes in local lipid composition and / or packing, coupled to a protein domain as has been documented by others^59–61^.

These results show that VASP forms clusters on both neutral and negatively charged membranes with Endophilin and Lamellipodin^850–1250^. The presence of Endophilin and VASP on GUVs leads to the formation of large protein domains that that likely promote the local formation of membrane tubules.

### Actin is enriched in, and polymerized in, Lamellipodin^850-1250^-VASP and Endophilin-VASP condensates resulting in a templated condensate reshaping

Protein condensates have been shown to act as reaction crucibles for various cellular processes^35–37^ including the polymerization of cytoskeletal proteins^34, 38–40^. Having identified the capacity of Lamellipodin^850–1250^-VASP and Endophilin-VASP to undergo LLPS (**Figure 2**), we probed the ability of the condensates to function as centers for actin polymerization.

We introduced actin (5 µM total, 5% Rhodamine green labeled) to preformed Lamellipodin^850-1250^(10 µM total, 2.5% Alexa 647 labeled)-VASP (20 µM total, 1.25% Alexa 594 labeled) condensates and observed a strong partitioning of actin into the condensates (∼70x, **Fig 3A**). From the images collected, we noticed a shift in the morphology of the condensates from spherical to donut-like structures enriched in VASP, Lamellipodin^850-1250^, and actin (**Fig 3A**). We stained the sample with Phalloidin (150 nM Alexa 647 labeled) and we observed ∼100x enrichment of Phalloidin in the condensates and we found strong colocalization between the actin and Phalloidin (**Fig 3B**). These results confirm that actin is enriched within Lamellipodin^850-1250^-VASP condensates, and that polymerization is localized within them.

**Figure 3:**
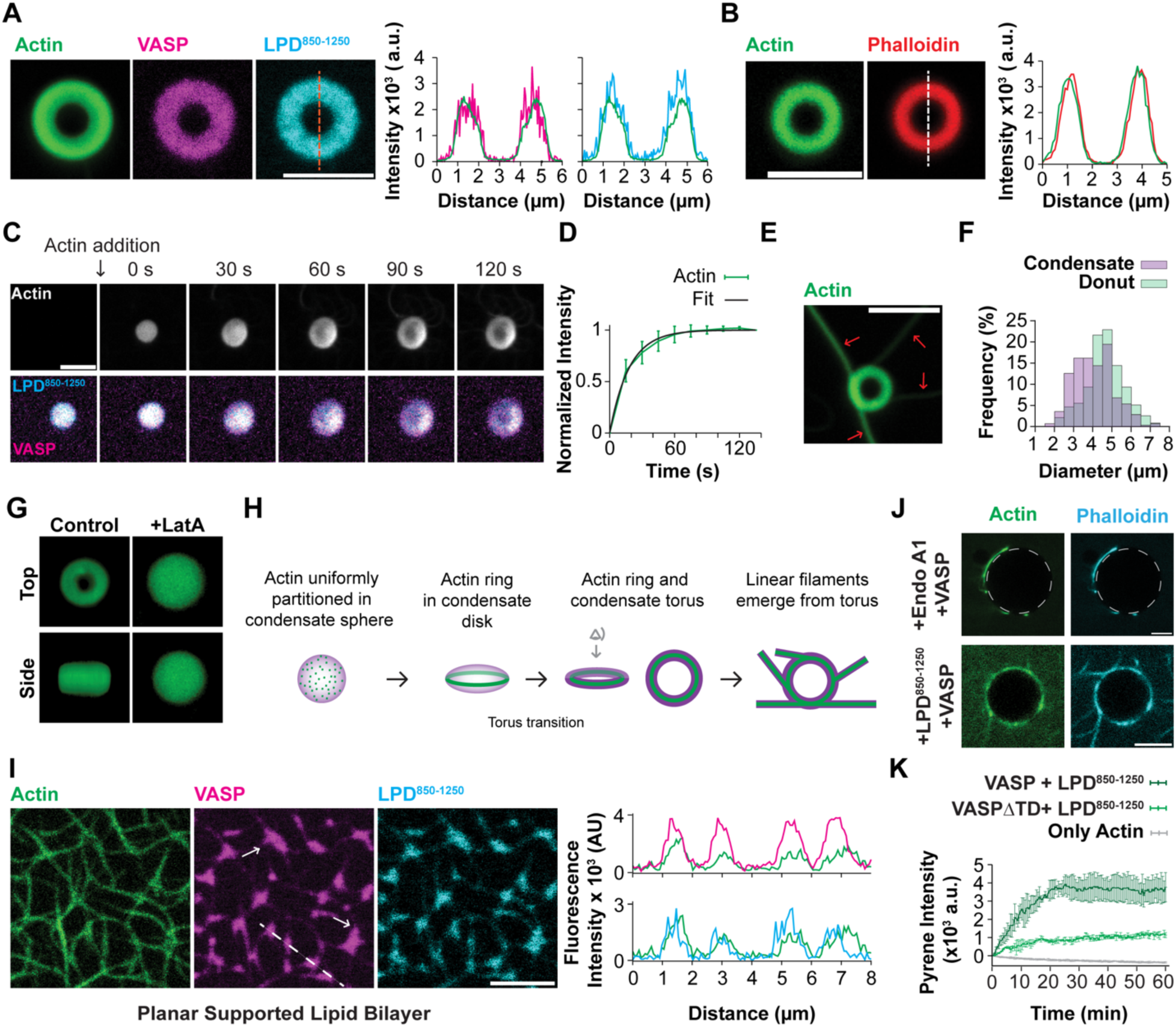
Actin polymerization from VASP containing condensates templates condensate deformation. **A** Actin (5 μM total with 5% Rhodamine green actin) polymerized within LPD^850-1250^ - VASP (10 μM, 20 μM respectively with 250 nM labeled protein each) condensates, imaged after 30 minutes of incubation. Scale bar 5 μm. Line intensity profile along dashed line shown in image on the left. **B)** Staining of actin polymerized in condensates prepared under conditions mentioned in A using Phalloidin Alexa 647. Line intensity profile along dashed line shown right of image. Scale bar 5 μm. **C)** Time evolution of actin (5 μM total with 5% Rhodamine green actin) polymerized in condensates formed with 20 μM VASP and 10 μM LPD^850-1250^. Images acquired in intervals of 30s show the time dependence of the peripheral distribution of actin and the concomitant deformation of the condensate. Scale bar 5 μm. **D**) Enrichment of actin in LPD^850-1250^-VASP condensates over time corresponding to conditions described in **C** (green curve with points). Mean fluorescence intensity was calculated for each time point and normalized with respect to mean intensity of the final time point. The normalized intensity profile was fit to a single exponential with a halftime of enrichment = 18.9 +/-7.1 s shown in the black curve. N = 11 condensates. **E)** Instance of actin polymerization under conditions described in A, resulting in the formation of linear filaments from the condensates (Red arrows point to linear filaments). Scale bar 5 μm. **F)** Size distribution of LPD^850-1250^-VASP condensates and actin rings formed upon polymerization of actin within condensates. Condensates of composition 20 μM VASP (1.25% Alexa 594 labeled protein) and 10 μM LPD^850-1250^ (2.5% Alexa 647 labeled protein) were incubated for 10 minutes after which actin (5 μM total with 5% Rhodamine green actin) or Actin buffer of equal volume was introduced and incubated for 30 minutes. Diameter of condensates and rings were measured manually using ImageJ. N condensates = 277 and actin rings = 236. **G)** Top and side views of a 3D reconstruction created from a Z-stack of actin polymerized in LPD^850-1250^-VASP condensates before (Left) and after treatment with Latrunculin A (Right). **I)** Schematic of confined actin polymerization and concomitant morphological changes in LPD^850-1250^-VASP condensates in the bulk. **I)** Left Actin (1 μM total with 25% Rhodamine green actin) polymerized on a SLB (1% Ni-NTA DOGS, 99% DOPC) with preformed His tagged LPD^850-1250^-VASP clusters (100 nM LPD^850-1250^ and 2.5 μM VASP), showing deformation of clusters with pointed exit points for actin (White arrows point to a few examples in both the VASP and LPD^850-1250^ channels). Imaged after 30 minutes of polymerization. Scale bar 5 μm. Right Line intensity profile corresponding to the dashed white line shown in Left plotted pairwise, left LPD^850-1250^ and Actin, right VASP and Actin. **J)** Representative images of actin polymerized (5 μM total with 5% Rhodamine green actin) on GUVs in the presence of Top (2μM) EndoA1 and VASP (1μM) or Bottom, LPD^850-1250^ (1μM) and VASP (1μM) stained with 150 nM Alexa 647 labeled phalloidin. The white dashed circle represents the location of the GUV. Scale bar 5 μm. **K)** Kinetics of actin polymerization with LPD^850-1250^ and VASP truncates was assessed using the pyrene fluorescence assay. LPD^850-1250^, VASP and VASP-ΔTD were kept constant at 1 μM each and actin was used at 2 μM unlabeled doped with 250 nM pyrene labeled actin. The proteins were diluted in the actin buffer (see methods), and actin was added immediately before the initiation of the experiment. The error bars represent the standard deviation of duplicates.

Using timelapse imaging, we captured the dynamics of actin polymerization and concomitant deformation of the condensate. From the time evolution series (**Fig 3C**), we observed a rapid enrichment of actin within the condensates (Top row, t = 0 s). Subsequently, we noticed the widening and flattening of the condensate (t = 30 s) into an oblate/disk shape. With further evolution, we noticed a depletion of Lamellipodin^850-1250^, VASP and actin from the center of the condensate with an enrichment of the three proteins at the periphery (t= 60-90s) forming a torus-like shape. We also observed the formation of linear filaments from the actin donuts (**Fig 3E**), suggesting that the donut shaped structures might be intermediates.

We quantified the enrichment dynamics of actin within the condensates and found that the halftime of enrichment was ∼ 18 s. We further noticed that the onset of the transition from disk to torus coincided with the saturation of actin fluorescence within the condensates (Fig **3C****, 3D** ∼ 60 s).

To test if the deformation of the condensate resulting in the formation actin rings was indeed due to the polymerization of actin, we asked if subsequent depolymerization of actin would return the condensate back to its initial spherical shape. We incubated the donut shaped actin polymers formed in Lamellipodin^850-1250^-VASP condensates (**Fig 3A)** with latrunculin-A (Lat-A), an actin depolymerizing drug. Indeed, upon depolymerization, we noticed a loss of the donut shaped actin structures, and observed a reemergence of spherical condensates, with a uniform actin fluorescence throughout the condensate, as verified by Z-stack imaging and 3D reconstruction (**Fig 3G**). We also found VASP and Lamellipodin^850-1250^ showed uniform fluorescence distribution upon the depolymerization of actin (**Supplementary figure 5**). These results conclusively show that enrichment of actin alone is insufficient, and polymerization is essential to deform the condensates to form donut shaped structures.

To identify additional factors that control the oblate-to-donut transition (**Fig 3I**), we turned our focus to the size of the condensates and the actin rings. We estimated the diameters of both the condensates and the actin rings (outer diameter), and we identified a shift in the size distribution of the actin rings to larger diameters compared to those of Lamellipodin^850-1250^ -VASP droplets (**Fig 3F, Supplementary** Figure 6). This size increase can be appreciated by following the actin channel in **Fig 3C**, which demonstrates a flattening and concomitant widening of the condensate within 60 s of the introduction of actin. This further supports the conclusion that condensate deformation is a consequence of actin polymerization within the droplets.

Ring-like actin structures have been previously observed in VASP condensates^40^, but our study is the first to report the formation of donut-like shaped condensate as a result of actin polymerization. A key difference in the experimental design employed in the two studies is the use of the molecular crowders. In contrast to the work of Graham, et. al^40^, we prepared VASP containing condensates in the absence of Polyethylene Glycol (PEG) and therefore suspect that a major factor influencing the condensate morphologies after actin polymerization is the presence of PEG. We and others have found that the presence of PEG can influence the mobility of proteins within condensate and cause solidification^23, 62^, including the study by Graham, et. al^40^. The possible increase in the stiffness of the condensate might prevent its widening upon actin polymerization, as observed in our system, consequently, resulting in the formation of actin rings, without the transition of the condensate to a torus. We conclude that the spherical shape of the condensate templates the formation of actin rings, which deform the condensate, and ultimately leads to a transition from spherical to toroidal topology (**Fig 3H**).

We wondered whether VASP rich condensates on membranes could behave like the bulk condensates in localizing and polymerizing actin. To address this question, we introduced actin (1 µM total, 25% rhodamine green labeled) to preformed Lamellipodin^850–1250^ -VASP (100 nM Alexa 647 labeled, 2.5 µM total 10% Alexa 594 labeled respectively) clusters on SLBs. We observed the enrichment of actin and the generation of filaments from the protein clusters resembling spikes-like projections (**Fig 3I, Supplementary figure 7**). We observed the membrane bound condensates reshape and colocalize with the actin filaments (**Fig 3I, white arrows**). Similarly, we observed the formation of filaments from Endophilin-VASP and Lamellipodin^850-1250^ -VASP clusters on GUVs (**Fig 3J**).

As multivalent interactions between VASP and Lamellipodin^850-1250^ are essential for their phase separation (Fig **2G**), we asked if the capacity to polymerize actin was also influenced by their multivalent binding. To test this, we monitored actin polymerization kinetics in the presence of Lamellipodin^850-1250^ with either VASP or VASP-ΔTD and found that indeed upon the loss of higher valency interactions, actin polymerization was hindered (Fig **3K**).

Collectively these results show that Endophilin-VASP and Lamellipodin^850-1250^ -VASP condensates can locally concentrate actin, and that actin polymerization is initiated within, and localized to, the condensates. This polymerization results in the deformation of protein droplets and clusters forming donut-like structures in the bulk and elongated spike-like features on the membrane.

### Actin to VASP and Endophilin ratios control actin morphology

We next asked how the concentration space for VASP, Endophilin and Lamellipodin^850-1250^ affects the final actin morphology. We generated Lamellipodin^850-1250^ and VASP condensates using protein concentrations between 5 and 20 µM each and polymerized actin in them (at a total solution concentration for actin of 5 µM). With increasing VASP:actin ratios, we noticed a larger population of actin ring like structures, with a lower abundance of linear filaments in the sample (morphologies that are predominantly linear are classified as linear in the text and in figure legend and captions) (**Fig 4A, 4B, Statistical analysis in Supplementary figure 8**). We observed a reduction of linear filaments from 88 % to 15 % with an increase in the VASP:actin ratio from 1:1 to 4:1. We also found that for a given VASP:actin ratio, varying the Lamellipodin^850-1250^:Actin ratio did not influence the morphological distribution (**Fig 4A, 4B, Supplementary figure 8**), suggesting that only VASP:actin ratios dictate actin filament morphology in Lamellipodin^850-1250^-VASP condensates.

**Figure 4:**
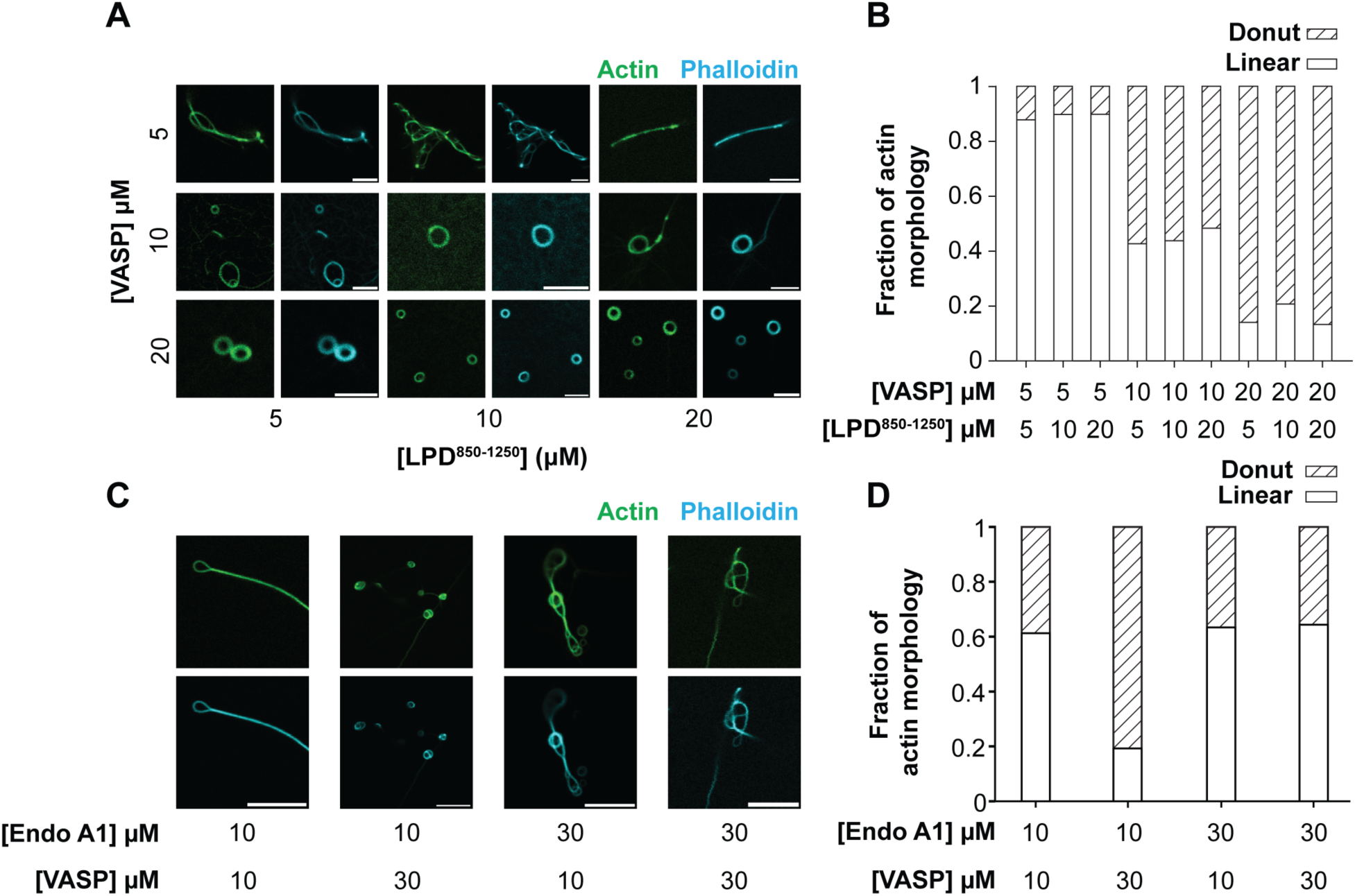
Endophilin-VASP and Actin-VASP ratios control actin morphology. **A** Actin (5 μM total with 5% Rhodamine green actin) polymerized in LPD^850-1250^-VASP condensates (5-20 µM each) stained with 150 nM Phalloidin-A647. Images representing the most frequent actin morphology observed for each composition of LPD^850-1250^ and VASP. Scale bar 5 µm. **B)** Distribution of actin morphologies (linear structures, in white, or donut-like structure, in dashed) observed for the conditions described in **A**. Left to right N =181,128, 148, 124, 89, 62, 57, 53, 361. **C)** Actin (1 μM total with 25% Rhodamine green actin) polymerized in Endo-VASP condensates (10-30 µM each) stained with Phalloidin-A647. Images representing most frequent actin morphology observed for each composition of EndoA1 and VASP. Scale bar 10 µm. **D)** Quantification of actin morphologies (linear structures, in white, or donut-like structure, in dashed) observed in conditions corresponding to those described in **C**. Left to right N = 207, 249 175, 76. All experiments performed in triplicates. Statistical analysis for **B, D** shown in **Supplementary** figure 8.

We next investigated the effect of the concentrations of Endophilin and VASP on the morphology of the polymerized actin. We prepared Endophilin-VASP condensates and polymerized actin (1 µM total, 25% rhodamine green actin) in them. At the lower Endophilin concentration (10 µM), we noticed a shift in the population of actin from less to more donut-like structures with increasing VASP’s concentration (**Fig 4C, 4D, Statistical analysis in Supplementary figure 8**). At the higher Endophilin concentration (30 µM), we did not observe a statistically significant shift in the frequencies of donut and linear actin filaments with varying the VASP concentration (**Supplementary figure 8**). These results show that Endophilin’s concentration with respect to VASP influences VASP mediated formation of actin donuts. We also noticed that although the VASP:actin ratio ranged between 10:1 and 30:1 (larger than those tested for VASP-Lamellipodin^850-1250^), we obtained a large fraction of linear filaments for all the conditions tested (**Fig 4C, 4D, Supplementary figure 8**). This result shows that unlike Lamellipodin^850-1250^, Endophilin influences actin morphology and promotes the formation of linear actin filaments.

These findings show that both Endophilin-VASP and Lamellipodin^850-1250^-VASP condensates localize and polymerize actin (**Supplementary figure 10**), and that the ratio of proteins used to form the condensates influence the final morphologies of actin polymers obtained.

### Actin polymerization is suppressed in Endophilin-VASP-Lamellipodin^850-1250^ condensates

Thus far, we have characterized actin polymerization in protein condensates comprised of VASP with either Endophilin or Lamellipodin^850-1250^. As all three proteins are known to colocalize in FEME, we next considered the ternary mixture comprised of Endophilin, Lamellipodin^850-1250^ and VASP. We generated condensates containing the three proteins in the bulk by incubating 25 µM of each protein (**Figure 5A, 5B**). To verify their liquid nature, we assessed the translational mobility of the proteins in the condensate using FRAP. From single exponential fits of the recovery profiles, we found that the proteins had high mobile fractions and quick diffusion kinetics, with mobile fractions of 80%, 93% and 70% and t_1/2_ of 8.48 s, 8.26 s and 9.06 s for Endophilin, VASP and Lamellipodin^850-1250^ respectively (**Fig 5B**) suggesting the formation of liquid-like condensates.

**Figure 5:**
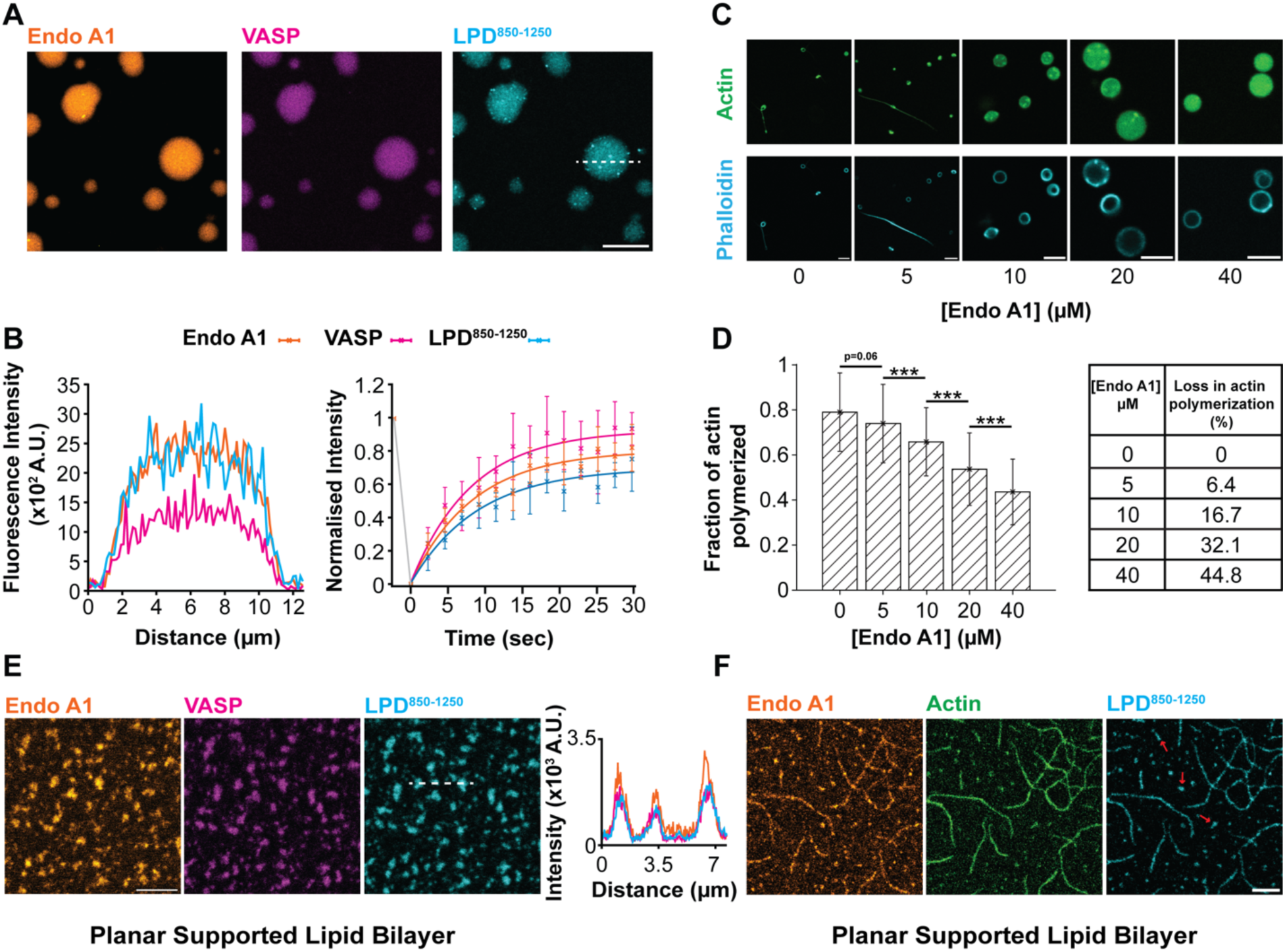
Endophilin-VASP-Lamellipodin^850-1250^ condensates have lowered actin polymerization efficacies. **A** Condensates created by the mixture of 25 μM each of EndoA1, VASP and LPD^850-1250^ doped with 2% EndoA1-Alexa 488, VASP-Alexa 594 and LPD^850-1250^ Alexa 647. Scale bar 10 μm. **B) Left** Line intensity profile along dashed line shown in **A,** showing colocalization of Endophilin, VASP and LPD^850-1250^. **Right** Normalized FRAP intensity curve of EndoA1, VASP and LPD^850-1250^ for conditions described in **A**. The error bars are the standard deviation of the intensities of 3 independent experiments. Solid lines represent single exponential fits of the average recovery. EndoA1 Mobile fraction 0.80, Halftime of recovery 8.48 s, VASP Mobile fraction 0.93, Halftime of recovery 8.26 s, LPD^850-1250^ Mobile fraction 0.70, Halftime of recovery 9.06 s. **C)** Actin (1 µM with 250 nM Rhodamine green Actin) polymerized in LPD-VASP condensates (10 µM each) in the presence of EndoA1 (0-40 µM), stained with 150 nM Phalloidin-A647. Images shown represent the most frequent actin morphology observed for each composition of EndoA1, LPD and VASP. Scale bar 5 µm. **D)** Quantification of the degree of actin polymerized for conditions shown in **C** calculated by the overlap of Phalloidin and rhodamine green intensity. Mean +/-S.D. shown in bar graph. Left to right n = 152, 762, 649, 960, 1194. Statistical analysis: Kruskal Wallis with post hoc Dunn test. *** corresponds to Bonferroni adjusted p value 0.0001.**E) Left** SLB (1% Ni-NTA DOGS, 99% DOPC) with preformed His tagged LPD^850-1250^ –VASP-Endophilin clusters (100 nM LPD^850-1250^ and 2.5 μM VASP, 2.5 µM Endophilin), showing colocalization of all proteins. **Right** Line intensity profile corresponding to the dashed white line shown in **Left.** Scale bar 5 µm. **Right** Line intensity profile across dashed line shown in **Left. F)** Actin (1 μM actin with 250 nM Rhodamine green Actin) polymerized in conditions described in **E**, imaged after 30 minutes of polymerization. Scale bar 5 μm.

We next asked if Endophilin-Lamellipodin^850-1250^-VASP condensates can localize and polymerize actin. We introduced actin (1 µM) to VASP-Endophilin-Lamellipodin^850-1250^ condensates (10 µM each) and subsequently stained them with phalloidin (Fig 5C middle panel). While we observed the enrichment of actin and positive phalloidin staining within the droplet, donut-shaped or linear filaments did not form (Fig 5C middle panel), suggesting weaker actin polymerization. Compared to the condensate systems investigated above (Fig **4**), in the three-protein condensates, we have newly introduced

Lamellipodin^850-1250^-Endophilin interactions, leading us to hypothesize that this interaction results in the lowered polymerization efficacy. To test this, we quantified the degree of actin (1 µM total, 25 % Rhodamine green labeled) polymerized, using Phalloidin (150 nM, Alexa 647 labeled), in a series of condensates with fixed VASP and Lamellipodin^850-1250^ concentrations (10 µM each), and variable Endophilin concentrations (0-40 µM) (**Fig 5C**). Consistent with our hypothesis, we observed a linear decrease in the colocalization of actin and Phalloidin (see methods for details) with increasing Endophilin concentrations suggesting an Endophilin dependent decrease in polymerization (**Fig 5C, 5D, Supplementary figure 9**). This result correlated with a shift in the actin morphologies from filaments and donuts (at low Endophilin concentrations) to spherical structures (at high Endophilin concentrations) resembling undeformed condensates (**Fig 5C**).

We then asked if VASP-Endophilin-Lamellipodin^850-1250^ condensates on lipid membranes also display a lowered capacity to polymerize actin? As VASP (**Fig 2H**) and Endophilin^23^ can individually form phase separated domains with Lamellipodin^850-1250^ on membranes, we hypothesized that Endophilin and VASP could co-phase separate with Lamellipodin^850-1250^ to form 3 protein condensates on lipid bilayers. To test this, we co-introduced Endophilin and VASP (2.5 µM each) to SLBs (99% DOPC/1% Ni-NTA DOGS) coupled with Lamellipodin^850-1250^ (100 nM) and found that the three proteins colocalized and formed phase separated clusters (**Fig 5E**). We added actin (1 µM total, 25% rhodamine green labeled) to the preformed clusters described above, and we observed the formation of sparse actin networks and puncta enriched in actin (**Fig 5F**). These results show that VASP-Endophilin-Lamellipodin^850-1250^ condensates polymerize actin to a lesser degree than VASP-Lamellipodin^850-1250^ condensates on membranes and compared to VASP-Lamellipodin^850-1250^ and VASP-Endophilin droplets in the bulk.

To further verify this conclusion, we quantified the extent of Lamellipodin^850-1250^-VASP mediated actin polymerization +/-Endophilin with the help of a pyrene fluorescence assay^63^. We monitored the polymerization of 2 µM Actin (with 250 nM Pyrene actin), in the presence of Lamellipodin^850-1250^ and VASP (1 µM each), +/-Endophilin (5 µM). From the fluorescence curves, we noticed a 75% decrease in the total actin polymerized in the presence of Endophilin (**Fig 6A, Supplementary figure 11**), providing additional evidence for the suppression of actin polymerization in the ternary protein system.

**Figure 6:**
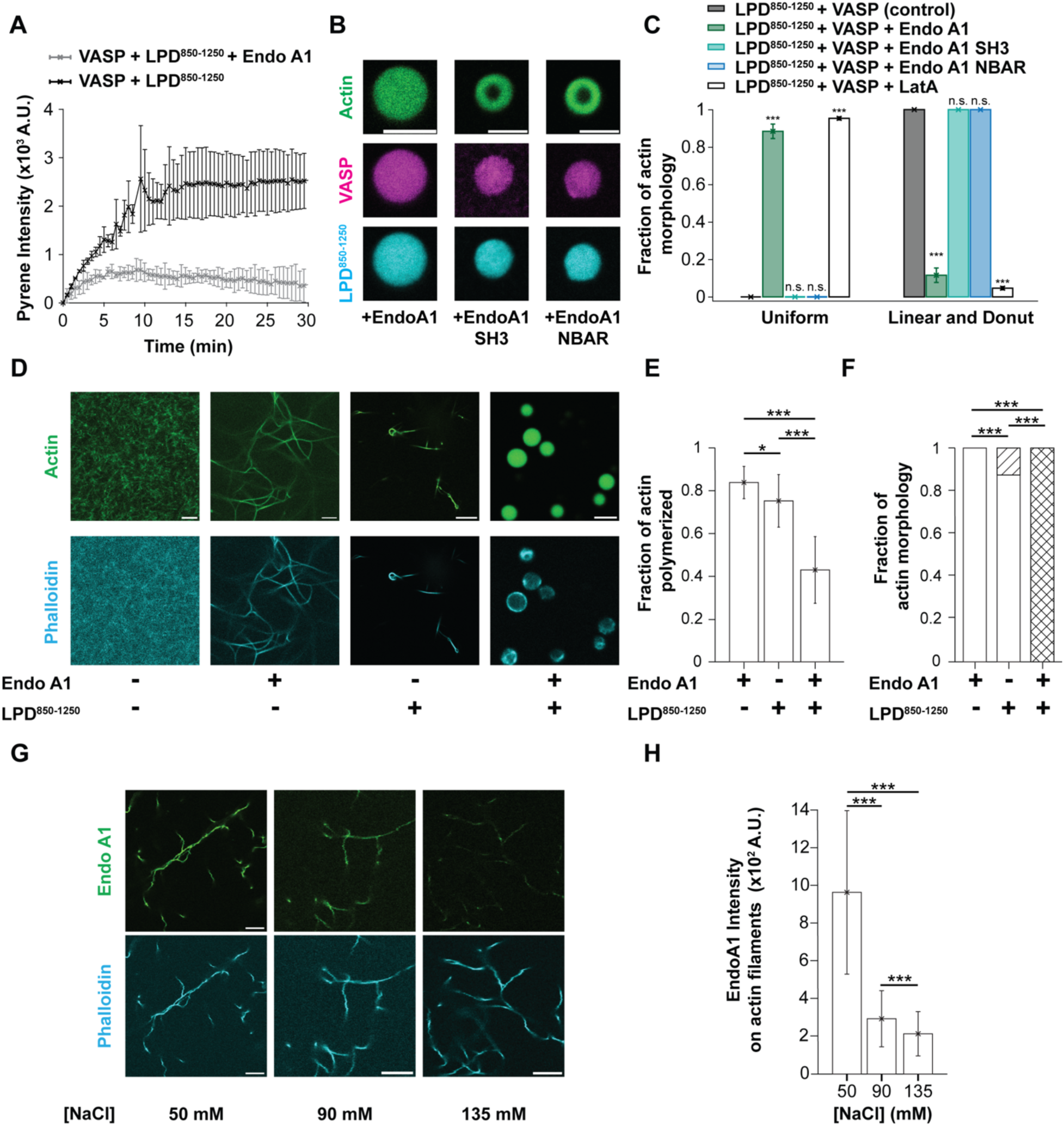
Endophilin condensates promote actin bundling; Endophilin-Lamellipodin interactions suppress actin polymerization. **A** Actin polymerization kinetics (2 μM actin + 250 nM Pyrene actin) in the presence of 1 μM LPD^850-1250^, 1 μM VASP +/-5 μM EndoA1. Error bars correspond to standard deviation of 3 independent measurements. **B)** Actin (5 μM total 5% Rhodamine green labeled) polymerized in condensates made of EndoA1 constructs (full-length EndoA1, monomeric SH3 and NBAR domains, VASP and LPD^850-1250^ (concentrations used were 40 μM EndoA1/SH3/NBAR, 20 μM VASP total 250 nM Alexa-594 labeled protein,10 μM LPD^850-1250^ total 250 nM Alexa-647 labeled protein). Images acquired after 30 minutes of polymerization. Scale Bar 5 μm. **C)** Percentages of actin morphologies observed after polymerization in condensates. Quantification of morphology of actin polymerized in 10 μM LPD^850-1250^ and 20 μM VASP condensates (grey bar, N =88), with 40 μM EndoA1 (green bar, N=76), SH3 monomer (teal bar, N=53), NBAR (blue bar, N=70), or 5 μM Lat-A (white bar, N = 66). Categorization was performed based on inspection of the actin distribution in the condensates. Statistical analysis: Chi-squared test, *** corresponds to Bonferroni adjusted p < 0.0001, n.s. corresponds to not significant. **D)** Actin (1 µM total, 25% Rhodamine green Actin) polymerized in EndoA1 or LPD^850-1250^ or EndoA1+ LPD^850-1250^ condensates (20 µM each protein, in the presence of 5 percent PEG), stained with Phalloidin A647. Scale bar 5 µm. Equatorial plane of the condensates shown in the EndoA1+ LPD^850-1250^ condition. **E)** Quantification extent of actin polymerized for conditions shown in **D**. Mean +/-S.D. shown in bar graph. Statistical analysis: Kruskal Wallis with post hoc Dunn test. *,*** correspond to Bonferroni adjusted p value <0.05 and < 0.0001. **F)** Distribution of morphology of polymerized actin corresponding to conditions shown in **D**. Left to right N = 205, 258, 355. Legend: Linear filament (white bar), donut (dashed lines) and uniform (crossed lines) actin morphologies. Statistical analysis: Chi-squared test, *** corresponds to Bonferroni adjusted p < 0.0001, n.s. corresponds to not significant. **G)**Interaction of Endophilin (250 nM A488 labeled EndoA1) with 100 nM polymerized Actin (stained with 150 nM Phalloidin-A647) at various salt concentrations (50-135 mM). Scale bar 10 µm. **H)** Quantification of intensity of EndoA1 on actin filaments corresponding to conditions described in **A**. Intensities from 3 independent trials used. Mean +/-S.D. shown in bar graph. Statistical analysis: Kruskal Wallis with post hoc Dunn test. *** corresponds to Bonferroni adjusted p value < 0.0001.**A,B** performed in 100 mM NaCl, 20 mM Hepes, 1 mM TCEP, pH 7.4 and 5% PEG. **G,H** performed in the following buffer: Variable NaCl, 20 mM Hepes, 1 mM TCEP, pH 7.4,

Revisiting the data shown in **Fig 5D**, with increasing concentrations of Endophilin, a larger fraction of Lamellipodin^850-1250^’s proline-rich motifs (PRMs, Endophilin binding sites) will be bound by Endophilin’s SH3 domain (mass action). This phenomenon is further promoted by Endophilin’s innate capacity to self-associate and form multivalent interactions with Lamellipodin^850-1250^ ^23^. As Endophilin and VASP bind distinct sites interspersed on Lamellipodin^850-1250^ ^17^ (**Supplementary figure 12**), with increased Endophilin binding, we would expect a decrease in the number of VASP molecules bound to Lamellipodin^850-1250^ due to hinderance/crowding. We would simultaneously expect limited VASP-Endophilin interactions due to competition for Endophilin’s SH3 domains between VASP and Lamellipodin^850-1250^. We hypothesize that these factors together reduce the clustering of VASP molecules on a molecular level, a key requirement for enhanced actin polymerization^12^, decreasing the degree of actin polymerized in VASP-Endophilin-Lamellipodin^850-1250^ condensates.

Multivalent Endophilin interactions (with itself and with Lamellipodin^850-1250^) are at the center of the above-described hypothesis. We therefore argued that the ternary protein system’s capacity to polymerize actin could be restored if multivalent Endophilin-Lamellipodin interactions were disrupted. To test this hypothesis, we replaced Endophilin with its individual domains (SH3 and NBAR) in the ternary system and prepared condensates of composition VASP-Lamellipodin^850-1250^ (20 µM and 10 µM respectively) and Endophilin-FL/ Endophilin-SH3/ Endophilin-NBAR (40 µM). We polymerized Actin (5 µM total, 5% Rhodamine green actin) within these condensates and investigated the degree of polymerization. Consistent with earlier findings, condensates containing full-length Endophilin showed uniform actin fluorescence indicating, at most, weak polymerization (**Fig 6B-Left, 6C dark green, Supplementary** Figure 13). However, when replaced by the individual domains (Endophilin-SH3 and Endophilin-NBAR), we observed the localization of actin to the periphery of the droplets indicating strong polymerization (**Fig 6B middle and right, 6C light green and blue**). Together, these findings show that only full length Endophilin can regulate the degree of actin polymerization, and that individual Endophilin domains (SH3 and NBAR) cannot impart this effect.

### Endophilin-Lamellipodin interactions regulate actin polymerization in condensates

Lamellipodin^850-1250^ contains 44 Arginine and Lysine amino acids, which are essential for its association with actin^12^. Along with binding actin, Lamellipodin^850-1250^ has been reported to promote actin bundling it in-vitro^12^, however factors regulating this function are not well established. We overlapped the location of Lamellipodin^850-1250^’s positively charged residues with its PRMs and found that 15 of the 44 basic residues (∼35%), are either within or flank the PRMs (**Supplementary figure 12**). We asked the question if the binding of Endophilin to Lamellipodin^850-1250^’s PRMs would modulate Lamellipodin^850-1250^’s capacity to bundle actin. To test this hypothesis, we polymerized actin (1µM total, 2.5% Rhodamine green labeled) in condensates containing Lamellipodin^850-1250^ (20 µM) or Endophilin-Lamellipodin^850-1250^ (20 µM each) (**Supplementary figure 14**) and quantified the degree of polymerization using Phalloidin staining (see methods). In agreement with our hypothesis, we observed a 50% decrease in the extent of actin polymerized for Endophilin+Lamellipodin^850-1250^ condensates compared to Lamellipodin^850-1250^ only condensates (**Fig 6D, 6E**). Consistent with our findings in the 3-protein condensates (**Fig 5C, 5D**), we observed a decrease in occurrences of linear filaments and donuts, with an increase in the population of spherical condensates with uniform actin fluorescence in the presence of Endophilin (**Fig 6F**). Together, these results provide evidence for a novel function of Endophilin as a promoter of actin bundling.

### Endophilin directly binds actin filaments and promotes actin bundling

The BAR proteins BIN1, PICK1 and pacsin2 all have recently been identified to bind actin^13-15^. We asked if Endophilin can bind actin directly. To address this, we tested the capacity of Endophilin (250 nM Alexa 488 labeled) to bind polymerized actin (100 nM actin, see methods). We visualized the polymerized actin by staining it with Alexa 647 Phalloidin and quantified the local Endophilin fluorescence on the filaments. We observed strong colocalization of Endophilin on actin filaments at 50 mM NaCl (**Fig 6G, 6H**). We noticed a decrease in the colocalization of Endophilin on the actin filaments upon increasing the ionic strength (from 50 mM to 135 mM), suggesting an electrostatically driven interaction.

As Endophilin can interact with both actin (**Fig 6G, 6H**) and itself^23^, we sought to understand if Endophilin could act as an actin cross-linker, promoting the formation of actin bundles. We introduced actin (1µM total, 2.5% Rhodamine green labeled) to Endophilin (20 µM) condensates and quantified the degree of polymerization using Phalloidin staining. Consistent with our hypothesis, we observed the formation of long bundles of actin filaments that colocalized strongly with Phalloidin (**Fig 6D-F**), providing evidence for a novel function for Endophilin as a promoter of actin bundling.

## Discussion

Recent studies have helped improve our understanding of the molecular mechanisms involved in the creation of endocytic priming patches and carriers in FEME^3, 4, 22, 64^. While actin is known to be indispensable, the initiators and regulators that control the extent of polymerization have been elusive. In this contribution, we show that VASP and Mena (members of the Ena/VASP family of actin polymerases) colocalize with FEME priming patches, and we identify a novel binding interaction between VASP and Endophilin. We establish noncanonical associations between the EVH1 (‘FPPPP’ motif recognizing) and EVH2 (actin binding) domains of VASP, with Endophilin’s SH3 domain. We further demonstrate that VASP forms liquid-like condensates in solution and on lipid membranes with the key FEME proteins Endophilin and Lamellipodin. We show that both bulk and membrane proximal condensates localize and polymerize actin, undergoing a templated shape transition. We reveal that the extent of polymerization in condensates can be controlled and modulated by multivalent Endophilin-Lamellipodin interactions. We finally show that Endophilin condensates can promote the polymerization and bundling of actin and we provide evidence for direct binding of Endophilin to polymerized actin.

### Model of actin polymerization and condensate deformation

From an energetic consideration, the cost of accommodating actin filaments within a condensate is the sum of the actin filament bending energy and the condensate interfacial energy, expressed as^40, 65^

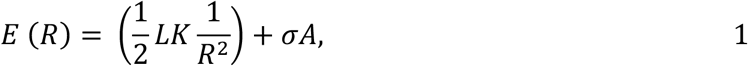

where 𝐸 is the total energy, 𝐾 is the bending stiffness of the actin filament, 𝐿 and 𝑅 are the length and radius of curvature of filament, respectively, 𝜎 is the surface tension, and 𝐴 is the surface area of the condensate.

In this work, we have shown that actin polymerization within condensates results in the deformation of the condensate following key shape transitions. The spherical condensate deforms to form a flattened oblate-like shape (Fig 3C-30 s, 3H) that subsequently transitions into a donut-like shape (Fig 3C-120 s, 3H). Here, we model the energies of actin filaments in a unit sphere condensate (radius = 1 µm) that undergoes the above-described shape transitions. We approximate the shape of the flattened condensate to a disk with curved edges and the donut shape to a torus (Fig 3H).

As actin filaments within condensates are known to align along the largest perimeter of condensates (Fig 3B-C)^40, 66^, for our model, we take the radius of curvature of the filaments (*R*) to be equal to the major radius of the condensate (radius corresponding to largest circumference of the condensate). From our experimental results, we find that upon the partitioning of actin into condensates, the volume of the condensate appears to be unchanged (Fig 3C). We therefore assume the total volume occupied by actin equal to 1% (negligibly small percentage) of the condensate volume and calculate the total number of actin monomers^67, 68^ and the number of filaments (*N*) in the condensate.

In Figure 7B, we plot the total energy of the actin/condensate system for a condensate undergoing deformation (increasing major radii) described by either a flattened condensate (disk with curved edges) or a torus of equal major radii. We use an experimentally determined value of the condensate surface tension, (10^-11^ N/µm)^69^ and a linearly scaled (by the number of actin filaments, *N*) value for the bending stiffness of actin^70^ to calculate the energy of the actin/condensate system (Fig 7A). We also plotted the difference in the energies of the two condensate shapes considered and noticed that for any radius larger than ∼1.6, a torus has a lower total energy than a flattened condensate (Fig 7B). This suggests that prior to the transition to a torus, a spherical condensate must undergo expansion (and concomitant flattening) which is consistent with our experimental findings (Fig 3F). To our knowledge, this is the first model aimed at capturing the deformation of condensates due to actin polymerization in 3 dimensions.

**Figure 7:**
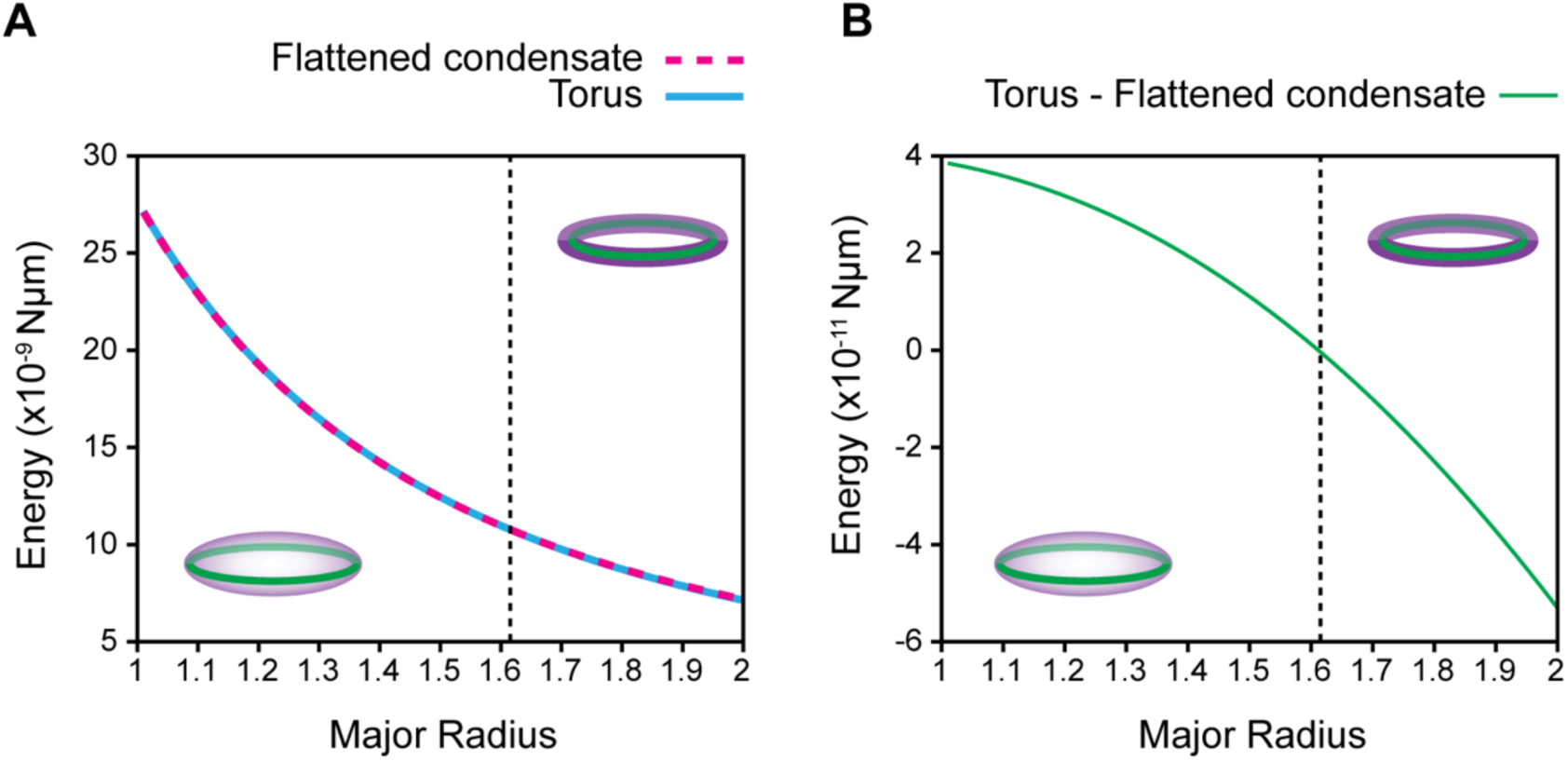
**A)** Plots representing total energy of a condensate (Eq 1) with actin filaments undergoing deformation at constant volume (unit sphere). Energy of the actin ring in a flattened condensate is shown in dashed magenta and in a torus in solid cyan. **B)** Plot showing the difference in energies of the two systems (actin ring in a torus or flattened condensate) considered. Dashed black line intersects at the point where the difference in the energies of the two shapes is equal to zero. Schematics depict the energetically preferred shape on either side of the dashed line.

### Biological consequences of findings Formation of FEME priming patches

It is known that local actin polymerization is required for the creation of priming sites in FEME^3^. While the role of actin in the generation and scission of endocytic buds has been reported on^6, 7, 71–73^, the need for actin polymerization in the development of priming patches is not well understood. Findings in this contribution and previous work by us^23, 56^ and others^12, 48^, have helped establish direct or indirect interactions between key FEME proteins including Endophilin, Lamellipodin and VASP with both actin and the lipid membrane. The innate capacity of these proteins to form liquid-like condensates coupled with the presence of interacting surfaces (actin and membrane) can lower the threshold concentration required to shift from a dilute to a condensed phase (known as prewetting)^74–77^. Collectively, we propose a model wherein the plasma membrane and proximal actin filaments act as preferential surfaces for the recruitment and clustering of key FEME proteins resulting in the formation of phase separated condensates. As actin networks near the plasma membrane are constantly reorganized^78, 79^, we predict that the local depolymerization of actin near FEME priming sites along with Cdc42-GAP activity^22^ would increase the threshold concentration required for phase separation resulting in a loss of the priming patch. This model explains the actin dependent formation and dissolution of the endocytic priming sites in FEME.

### Promoting membrane scission

FEME carriers colocalize with numerous proteins including Endophilin^3^, however neither Lamellipodin^3^ nor VASP (**Fig 1B, 1C**) are found on them, implying the presence of a protein sorting step prior to scission. We propose that a functional consequence of the sorting step is the initiation of local actin polymerization promoting membrane scission. Assuming that the concentration of Endophilin, Lamellipodin and VASP in a FEME priming site are relatively fixed, an endocytic bud formed from this site would reduce the effective Endophilin concentration at the base where Lamellipodin and VASP are present. As a reduction in Endophilin-Lamellipodin interactions promotes actin polymerization (Fig 6A-C), we would expect increased polymerization at the base of the bud promoting membrane scission by the edge pushing model^71^.

### Summary

In conclusion, in this study, we identify the presence of Ena/VASP family of actin polymerases at FEME priming sites. We show that VASP forms liquid-like condensates with Endophilin and Lamellipodin, both in the bulk and on lipid membranes, and that the condensates function to localize actin polymerization. We provide evidence for novel actin binding and bundling activities of Endophilin and establish a function for Lamellipodin-Endophilin complexes as modulators of actin polymerization. We finally propose a model that explains the role of actin reorganization in the genesis and dissolution of endocytic priming patches and describes the spatiotemporal regulation of actin polymerization in FEME.

## Materials and Methods

### Reagents

Fluorophores used were Alexa Fluor™−488 C5-maleimide, Alexa Fluor™−488 5-SDP ester, Alexa Fluor™ 594 succinimidyl ester and Alexa Fluor™647 succinimidyl ester and were purchased from Invitrogen. Latrunculin A (Cayman) Alexa Fluor™ 647 Phalloidin (Invitrogen) were used in the actin depolymerization and staining respectively. Pyrene actin was purchased from Cytoskeleton.

Lipids used namely, 1,2-dioleoyl-sn-glycero-3-phosphocholine (DOPC), 1,2-dioleoyl-sn-glycero-3-phosphoethanolamine (DOPE), 1,2-dioleoyl-sn-glycero-3-phospho-L-serine (DOPS) and 1,2-dioleoyl-sn-glycero-3-[(N-(5-amino-1-carboxypentyl) iminodiacetic acid) succinyl] (nickel salt) (Ni-NTA DOGS) were acquired from Avanti Polar lipids (USA). Fluorescent lipids used in the study were Marina Blue™ 1,2-Dihexadecanoyl-sn-Glycero-3-Phosphoethanolamine (Marina Blue™ DHPE) (Invitrogen) and Forte Blue DHPE (AAT Bioquest).

HEPES, NaCl, Na_2_HPO4, NaH_2_PO_4_, DTT, IPTG, glutathione, EDTA were purchased from Fisher Scientific. Glycerol and ATP-Mg, Polyethlene glycol (avg. MW 8000) were purchased from Sigma Aldrich, Imidazole from Acros organics and Pierce™ TCEP-HCl from Thermo Scientific. Passivating agents used were Bovine serum albumin (BSA) (Fatty acid free, fraction V) (MP Biomedicals) and β-Casein (Thermo Fisher Scientific). PEG-PLL was a kind gift from Elizabeth Rhoades. Restriction enzymes and T7 ligase were purchased from New England Biolabs.

### Plasmids

#### Bacterial expression plasmids

Sequences coding for Lamellipodin^850-1250^, VASP, and Rat Endo A2 were synthesized (Biomatik) and inserted into PGEX-6P1 vectors, whereas the Endo A1 linker (SH3)5 was cloned into a PMAL vector with an N and C terminal TEV cleavage site for the MBP and 6xHis tags respectively. VASP-ΔTD (AA 1-335), and EVH1 (AA 1-113) were cloned using the VASP plasmid as the template. DNA encoding for Rat Endo A1 (single cysteine mutant) (Endo-A1), SH3, NBAR, ΔSH3 were inserted into the PGEX-6P1 vector, and its generation is described elsewhere^80^.

#### Mammalian expression plasmids

VASP (pEGFP-C1-VASP) was a kind gift from Tatyana Svitkina. VASP and its truncates EVH1 (AA 1-113), linker (AA 114-224) and EVH2 (225-380) were cloned into the mScarlet vector.

### Confocal Microscopy

An Olympus IX83 inverted microscope equipped with a 60x 1.2 NA water immersion objective (Olympus) and a FluoView 3000 scanning system (Olympus, Center Valley, PA) was employed for confocal microscopy and FRAP studies. Sequential line scanning was implemented when imaging samples with multiple fluorophores (Marina blue, Alexa 488, Alexa 594, and Alexa 647) to cross-talk between channels. Images were analysed using Fiji and python (v 3.7.1).

For condensate and actin polymerization studies, coverslips (25 mm, Fisher Scientific) were cleaned by bath sonication in 2% Hellmanex, followed by three washes with water. The cleaned coverslips were either used immediately or stored in 200 proof ethanol until they were needed. The coverslips were passivated by creating a PEG-PLL coating following the protocol described elsewhere^81^. The sample to be imaged was deposited in the centre of the coverslip and a closed imaging chamber was created by sandwiching another coverslip on top with a thin layer of vacuum grease (Molykote, Dupont) around the edge.

### BSC1 cell culture

BSC1 cells were cultured at 37°C 10% CO2 in DMEM (Sigma) supplemented with 10% FCS and Penicillin / Streptomycin. Cells were plated on uncoated glass coverslips for immunofluorescence experiments at a density of approximately 2.5×10^4^ cells per coverslip.

### HeLa culture and lysate preparation

HeLa cells were cultured in DMEM supplemented with 10% fetal bovine serum (FBS), and 1x Penicillin-Streptomycin antibiotic. For generating cell lysate, the cells were seeded on 100 mm dishes and transfected with 2-3 µg of DNA at 80% confluency. Cells were harvested in cold DPBS after 24 h of over-expression and lysed by tip sonication.

### Immunofluorescence

Cells were left to spread on glass coverslips for 6 hours and were fixed with 4% PFA in PHEM buffer (60mM PIPES, 25mM HEPES, 10mM EGTA, 2mM MgCl_2_.6H_2_O, 0.12M sucrose) for 20 minutes at 37°C. Fixed cells were washed three times with cytoskeletal TBS buffer (cTBS: 10x stock comprised of 200mM Tris-HCl, 1.54M NaCl, 20mM EGTA, 20mM MgCl2 x 6H2O, pH7.5). Fixed cells were quenched for 10 mins with cTBS containing 50mM NH_4_Cl before being blocked and permeabilised for 1 hour at room temperature with cTBS containing 10% BSA, 10% normal goat serum and 0.05% saponin. Cells were then washed once with cTBS. Cells were stained with primary antibodies overnight at 4°C followed by secondary antibodies for 1 hour at room temperature. Antibodies were used at the following dilutions in cTBS with 1% BSA and 0.05% saponin: anti-Endo A2 1:200 (mouse monoclonal sc-365704, Santa Cruz Biotechnology), anti-VASP 1:400 (rabbit polyclonal)^82^, anti-Mena 1:400 (rabbit polyclonal)^82^, Alexa 488 goat anti-mouse and Alexa568 goat anti-rabbit 1:500 (Invitrogen). Cells were washed three times with cTBS after both primary and secondary antibody incubations. Cells were mounted overnight at room temperature using Prolong Diamond mounting reagents (Invitrogen). Cells were imaged using a CSU-W1 SoRa system (Nikon) with a 60x objective and an intermediate magnification of 2.8x.

### Protein expression and purification

BL21-CodonPlus(DE3)-RIL cells were transformed with the plasmid expressing the protein of interest. Single colonies were picked for overnight starter culture growths. Large volume growths were initiated using 25 mL of starter culture per litre of media and grown to an O.D. 600 of ∼ 0.7. Cultures were cooled and induced at 18 °C overnight with IPTG (0.3 mM for Lamellipodin^850-1250^, VASP WT, VASP-ΔTD, EVH1 and GST-Endo A1 ΔSH3, and 0.6 mM for Endo A1, SH3, NBAR. Cultures were spun down at 4k rpm at 4 ° C, resuspended in a lysis buffer (check specific protein) with 1 mM PMSF. The cell suspension was lysed using a tip sonicator followed by centrifugation at 15k rpm for 1 hour to separate the lysate from insoluble cellular debris. The lysate was filtered with a 0.22 µm syringe filter (Millipore-sigma) prior to affinity chromatography.

### The buffers used in the purifications are as follows

GST lysis buffer: 300 mM NaCl, 50 mM Tris, 2 mM DTT, 1 mM EDTA, pH 8 GST elution buffer: 150 mM NaCl, 50 mM Tris, 2 mM DTT, 1 mM EDTA, 20 mM glutathione, pH 8.

His lysis buffer: 300 mM NaCl, 20 mM Imidazole, 2 mM DTT, pH 8

His elution buffer: 150 mM NaCl, 50 mM Tris, 500 mM Imidazole, 2 mM DTT, pH 8

Cation exchange A buffer (SA): 150 mM NaCl, 20 mM Sodium Phosphate, 1 mM DTT pH 7

Cation exchange B buffer (SB): 1 M NaCl, 20 mM Sodium Phosphate, 1 mM DTT pH 7 Anion exchange A buffer (QA): 150 mM NaCl, 50 mM Tris, 1mM EDTA, 1 mM DTT pH 8 Anion exchange B buffer (QB): 1 M NaCl, 50 mM Tris, 1mM EDTA, 1 mM DTT pH 8 General storage buffer (S buffer): 150 mM NaCl, 20 mM Hepes, 1 mM TCEP, pH 7.4.

VASP storage buffer: 200 mM NaCl, 25 mM Hepes, 1 mM TCEP, 5% glycerol, pH 7.5

### Lamellipodin^850-1250^

The lysate was incubated with Ni-NTA agarose beads (Gold Bio) overnight at 4 °C. The beads were washed extensively with the His lysis buffer and incubated with PreScission protease overnight to cleave the GST tag. The beads were further washed with the lysis buffer to remove free GST, and the protein was eluted with the His elution buffer. The eluted protein was buffer exchanged using a HiTrap desalting column into the SA buffer. The protein was further purified using a HiTrap SP HP cation exchange column on a NaCl gradient using the SA and SB buffers. Pure protein fractions were confirmed using SDS-PAGE and were passed on a Superdex S200 pg 16 column (Cytiva) equilibrated with the S buffer.

### VASP WT, VASP-ΔTD, EVH1

Ecoli expressing the above mentioned VASP constructs were lysed in the GST lysis buffer with 5% glycerol and with PMSF and DNAse. Sepharose 4B glutathione beads were added to the filtered cell lysate and incubated overnight at 4 °C. The beads were washed extensively with the lysis buffer and subsequently incubated with PreScission protease overnight at 4 °C. The beads were spun down and the supernatant containing the protein of interest was isolated. The protein was concentrated and passed on a Superdex S200 column equilibrated with VASP storage buffer. The fractions corresponding to pure VASP were identified using SDS-PAGE.

### Endo A1, EndoA1 SH3, Endo A1 NBAR, GST-Endo A1, GST-Endo A2, GST-Endo A1 ΔSH3

Endo and its subconstructs were lysed using the GST lysis buffer. The clarified lysate was applied to a GSTrap^TM^FF column (GE Healthcare) and washed extensively with the lysis buffer. The GST tagged protein was eluted using the GST elution buffer containing glutathione. The GST tag was cleaved using PreScission overnight at 4 °C. The GST tag was separated from the protein of interest using an anion exchange chromatography step. The fractions post the ion exchange that contained pure protein was determined using SDS-PAGE and were concentrated and passed on a Superdex S200 pg 16 column (Cytiva) as the final step of the purification. For the GST tagged Endo constructs, the purification after the elution from the GST column continued with the size exclusion chromatography step.

### Actin

Actin was a kind gift from Elizabeth Rhoades. It was purified based on the method described here^83^.

### Protein labelling

The VASP constructs were labelled N terminally using Alexa Fluor 594 SDP ester. Lamellipodin^850-1250^ was labelled using Alexa 647 succinimidyl ester. Endo was labelled using a C5-Maleimide Alexa 594 fluorophore. The labels were added at 7-9-fold excess. The reactions were performed in their respective storage buffer and allowed to occur overnight (16-20 h). The labelled protein was separated from free dye by passing the sample over two serially attached HiTrapTM (5 mL) desalting column.

### Protein condensate formation and actin polymerization in the bulk

The experiments were performed in the S buffer with the desired ionic strength. Given the varying salt contributions from the proteins (refer protein storage buffers), the overall ionic strength was adjusted with 20 mM Hepes pH 7.4 and 1 mM TCEP to obtain the desired final concentration. The protein mixtures were incubated for 10 min at room temperature (21-23 °C) prior to imaging or addition of actin. For fixed time actin polymerization assays, a mixture of unlabelled (4.75 µM) and Rhodamine-green (250 nM) labelled actin was added to the condensate mixture and allowed to polymerize for 30 min at room temperature with a final ATP concentration of 25 µM in the solution. The samples were imaged on PEG-PLL passivated chambers.

For generating the size distribution histogram, condensates were first formed in 120 mM NaCl as described above and allowed to incubate for 10 minutes. Actin (5 µM) or Actin buffer of equal volume was added to the condensates and allowed to evolve for 30 minutes. The samples were deposited on PEG-PLL passivated closed chambers and imaged using confocal microscopy. The size of the condensates and actin rings was measured manually using ImageJ. The data was binned in 0.5 µm intervals.

### Actin polymer staining and actin depolymerization

Alexa647-Phalloidin was used to stain polymerized actin. It was added at a final concentration of 150 nM at the end of actin polymerization and were allowed to incubate with the sample for 30 min at room temperature. Samples were subsequently imaged.

Latrunculin A (Lat-A) was used at a final concentration of 5 µM. It added at the end of actin polymerization and was allowed to incubate with the sample for 30 min at room temperature.

### Preparation of supported lipid bilayers (SLB)

Lipid stocks composed of 98:1:1 mol percent DOPC: Ni-NTA DOGS: Marina Blue DHPE lipids were prepared at 5 mg/mL in chloroform. A desired volume of lipids was dried under nitrogen gas and dried further under vacuum between 2 and 16 h. Liposomes were resuspended in S buffer at a concentration of 0.2 mg/mL by vortexing. The suspension was sonicated in a water bath for 30-40 min and subjected to 5 freeze-thaw cycles. The vesicle suspension was finally extruded 13-17 times through a polycarbonate membrane with a pore size of 100 nm. The LUVs were used immediately for preparation of the SLBs.

8 well chambered coverslips (Cellvis) were used for the preparation of SLBs. The wells were first rinsed with 50% (v/v) isopropyl alcohol followed by Milli-Q water. 10 M NaOH was added into the wells for an incubation of 2 h. The wells were thoroughly rinsed with Milli Q water. LUVs were added to the cleaned chambers (150 µL, 0.2 mg/mL) for 30 min. To dislodge unruptured LUVs, the SLBs were rinsed with 650 µL of S buffer thrice. During each rinse, the buffer was repeatedly passed on top of the bilayer without introducing air or scratching the surface, with a minimum volume of 150 µL always maintained in the well. The bilayers were treated with BSA (final concentration 1 mg/mL) for 30 min to passivate any defects. The wells were rinsed thrice with S buffer and were ready for protein conjugation. The quality of the SLB was verified by performing FRAP.

### Coupling proteins to SLBs

Poly histidine tagged Lamellipodin^850-1250^ was coupled to SLBs by adding 100 nM total protein (10-20 % labelled protein) to a well. The protein was allowed to bind Ni-NTA functionalised lipids for 15 min, after which unbound protein was removed by gently rinsing the SLBs twice to achieve a 16x dilution. The protein-bilayer system was allowed to relax for 30 min prior to further experiments.

### VASP interaction with Lamellipodin^850-1250^ coupled SLB and actin polymerization

VASP (2.5 µM total) was added to the bilayers with the labelled protein concentration of 250 nM. The SLBs were imaged after 30 min of interactions. Actin (1 µM unlabelled with 100 nM Rhodamine-green labelled) was added at the end of the incubation and images were acquired after 30 min of polymerization.

### Correlation analysis

We performed radially averaged normalised auto- and cross-correlation analysis based on a method described elsewhere^23^.

### GST pull-down assay

Speharose 4B glutathione beads (Cytiva/GE) stored in 20% ethanol were spun down (500-1000 rpm) and washed in water followed by the S buffer. GST tagged protein was coupled to the beads by incubating 25 µM protein with beads at room temperature for 15 min. The beads are spun down and washed twice with the same buffer to ensure removal of uncoupled protein.

The conjugated beads are incubated with mScarlet VASP (and domains) expressing cell lysate or purified VASP or EVH1 (5 µM with 250 nM Alexa Fluor labelled protein) for one hour at room temperature. The beads were washed twice with buffer and imaged.

The fluorescence intensity on the beads were analysed using Image-J. A manual region of interest is drawn on the perimeter of the bead. The integrated intensity in the ROI is corrected for background fluorescence. The fluorescence data plotted in Fig 1 for is normalised by the mean value of the background corrected intensity of the corresponding fluorescent protein pulled down by GST-Endo ΔSH3.

A pull-down value greater than 1 is classified as SH3 mediated interaction and a value around or less than 1 is not considered to be an SH3 domain mediated binding.

### Pyrene actin polymerization assay

Actin polymerization kinetics was assessed by monitoring pyrene labelled actin (Cytoskeleton Cat #AP05) fluorescence. The polymerization was performed in the Actin buffer (5 mM Tris, pH 8.0 and 0.2 mM CaCl2) supplemented with 0.2 mM ATP and 0.5 mM DTT. The final concentrations used were 2 mM Actin with 250 nM pyrene actin, with VASP constructs and Lamellipodin^850-1250^ were at 1 µM, Endo and (SH3)_5_ were used maintaining the SH3 monomer concentration fixed at 5 µM. Measurements were performed on a Tecan M1000 plate reader.

### Giant Unilamellar vesicle experiments

GUVs of lipid composition DOPC/DOPE/marinaBlue-DHPE (69:30:1 mole percent) DOPC/DOPS/DOPE/MarinaBlue-DHPE (24:45:30:1 mole percent) were prepared using electroformation method as described elsewhere^84^. Briefly, 50 μl of the lipid mixture in chloroform (5 mg/ml) was added drop wise onto indium tin oxide (ITO) coated slides and spin coated at 1000 rpm for 1 min. In order to ensure the complete removal of residual solvent, the slides were dried under vacuum for at least 2 h. The lipid films were gently hydrated with aqueous 300 mM sucrose solution and electroformation was carried out at room temperature. An alternating current of amplitude 4 V peak to peak, and frequency 4 Hz was applied across two ITO coated slides for 2 h. The GUVs obtained were stored at room temperature and used after overnight resting.

Experiments involving the recruitment of VASP, polymerization of actin on GUVs and staining with phalloidin was performed as follows. The proteins and small molecules used in the experiment were first diluted in an osmotically balanced buffer (with respect to the sucrose solution used for GUV preparation) after which the GUVs were added to the mixture. The GUVs imaged immediately using a closed coverslip passivated with β-Casein.

### Preparation of polymerized actin

5X Polymerization buffer: 50mM Mops, pH7.0, 125mM KCl, 5mM EGTA, 5mM MgCl2 Actin was polymerized by incubating 1 µM actin with 1X polymerization buffer supplemented with 1 mM DTT and 2 mM ATP for 1 hour at room temperature.

### Binding of Endophilin to actin filaments

250 nM Alexa 488 labelled EndoA1 was incubated with 100 nM polymerized actin and 150 nM Phalloidin-A647 for 15 minutes at room temperature in 20 mM Hepes pH 7.4, 1 mM TCEP, and varying amounts of NaCl (50-135 mM). Samples were subsequently imaged on PEG-PLL coated coverslips.

### Analysis to estimate fraction actin polymerized within condensates

In these experiments, 1 or 5 µM (depending on the experiment) monomeric actin with 250 nM rhodamine green labelled actin was polymerized in presence of EndoA1-VASP, Lamellipodin^850-1250^ -VASP, EndoA1-Lamellipodin^850-1250^ or EndoA1-Lamellipodin^850-1250^ - VASP condensates for 10 minutes and stained with phalloidin-A647 (150-300 nM) for 10 minutes to visualize the polymerized actin.

To calculate the conversion efficiency of monomer to polymerized actin, an overlap percentage of polymerized:monomer actin was calculated for each structure identified. The following is the workflow.

Images acquired had labelled actin and phalloidin allowing for simultaneous visualization of monomeric and polymerized actin. Actin features/structures were isolated in the monomer actin channel using contour identification implemented in OpenCV. For the region within the actin contour, thresholding was performed in both the actin and phalloidin channels. The extent of overlap of the thresholded images was estimated giving the conversion percentage.

## Acknowledgements

This research was financially supported by the National Institutes of Health, grant NO GM 097552.

## Notes

### Competing Interest Statement

The authors have declared no competing interest.

### Summary of Updates

All the figures and the majority of the text has been revised.

