## Supplementary Figures for "Endophilin-Lamellipodin-VASP, key components in fast endophilin-mediated endocytosis, control actin polymerization within liquid-like condensates"

Supplementary figures and text

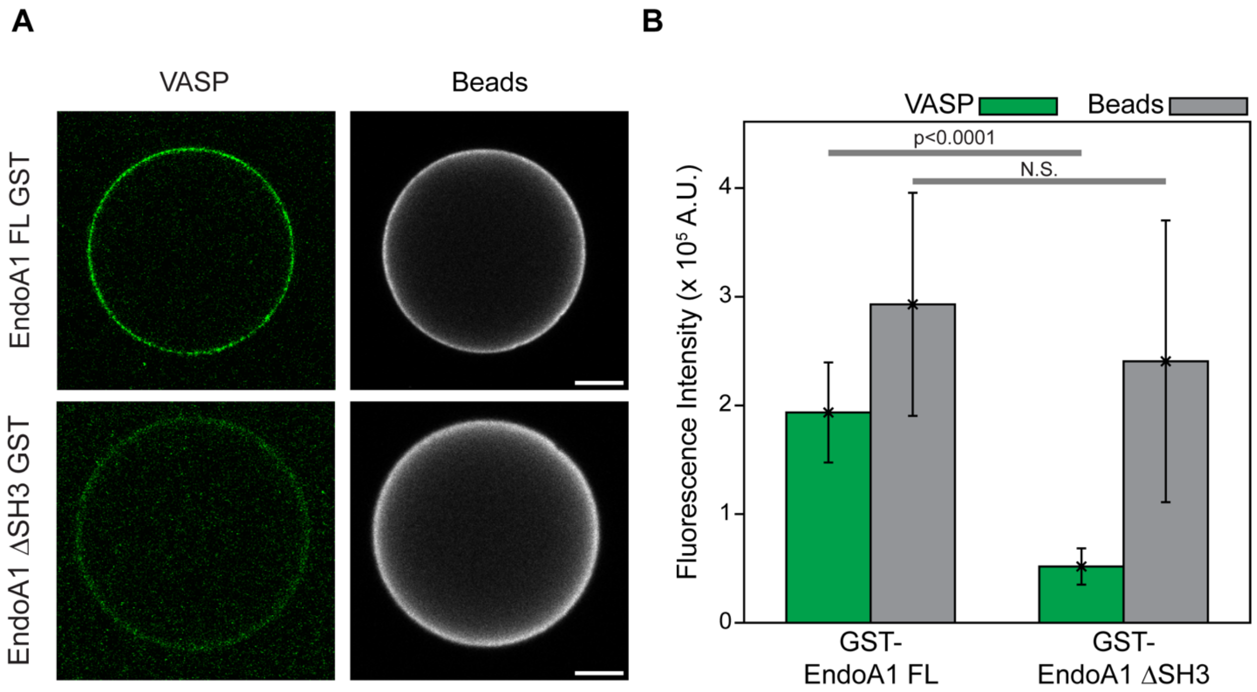


**Supplementary Figure 1: Assessment of coupling of GST tagged EndoA1 and EndoA1-ΔSH3 to Glutathione Sepharose 4B beads and subsequent pull down of eGFP-VASP from HeLa cell lysate.**

A) Representative images of eGFP VASP (green) and Alexa 647 labeled GST tagged EndoA1 and EndoA1-ΔSH3 (500 nM) doped with 25 μM unlabeled GST tagged EndoA1 and EndoA1-ΔSH3 coupled to glutathione beads. Scale Bar 20 μm.

B) Quantification of the fluorescence intensity of the eGFP and Alexa 647 coupled GST tagged protein on glutathione beads showing no significant difference in the coupling of GST tagged protein to the beads. Statistical analysis performed was an unpaired 2-tailed t test performed.

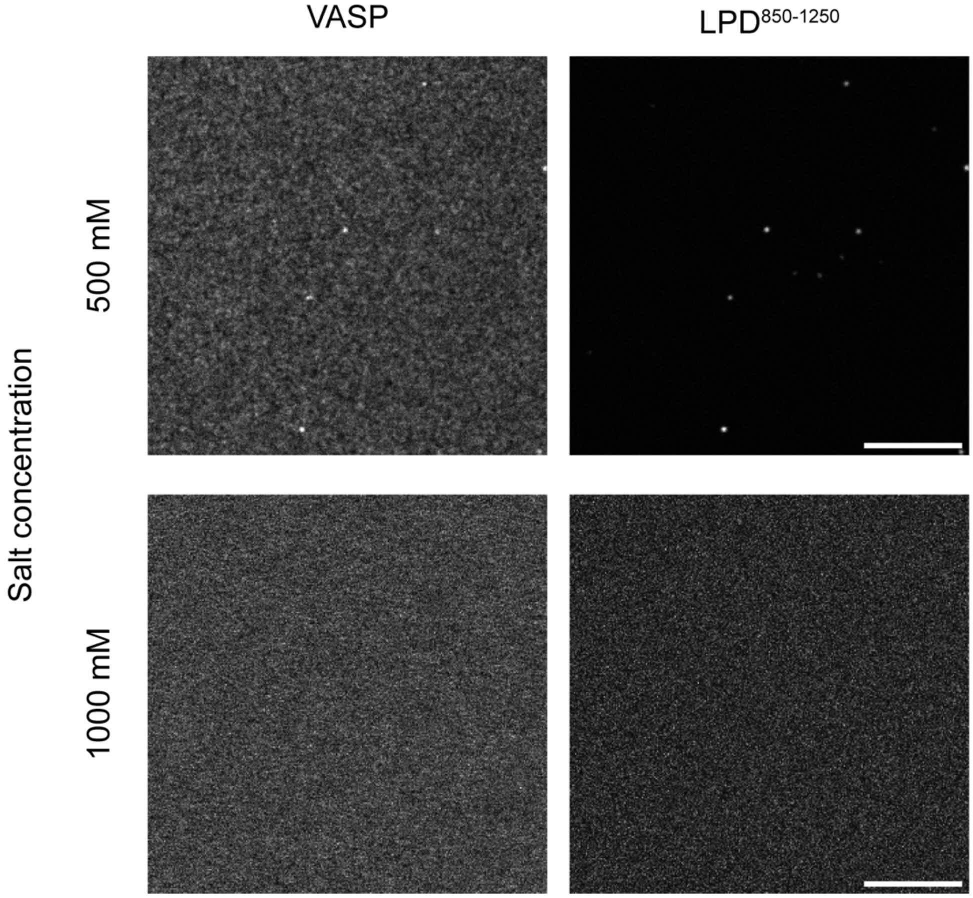


**Supplementary Figure 2: Effect of ionic strength on the phase separation of LPD$^{850\text{-}1250}$ and VASP**

20 µM VASP and 10 µM LPD$^{850\text{-}1250}$ doped with Alexa 594 and 647 labeled VASP and LPD$^{850\text{-}1250}$ in the presence of 500 mM (**Top row**) and 1 M (**Bottom row**) NaCl, imaged after 10 minutes of incubation at room temperature. Scale Bar 10 µm.

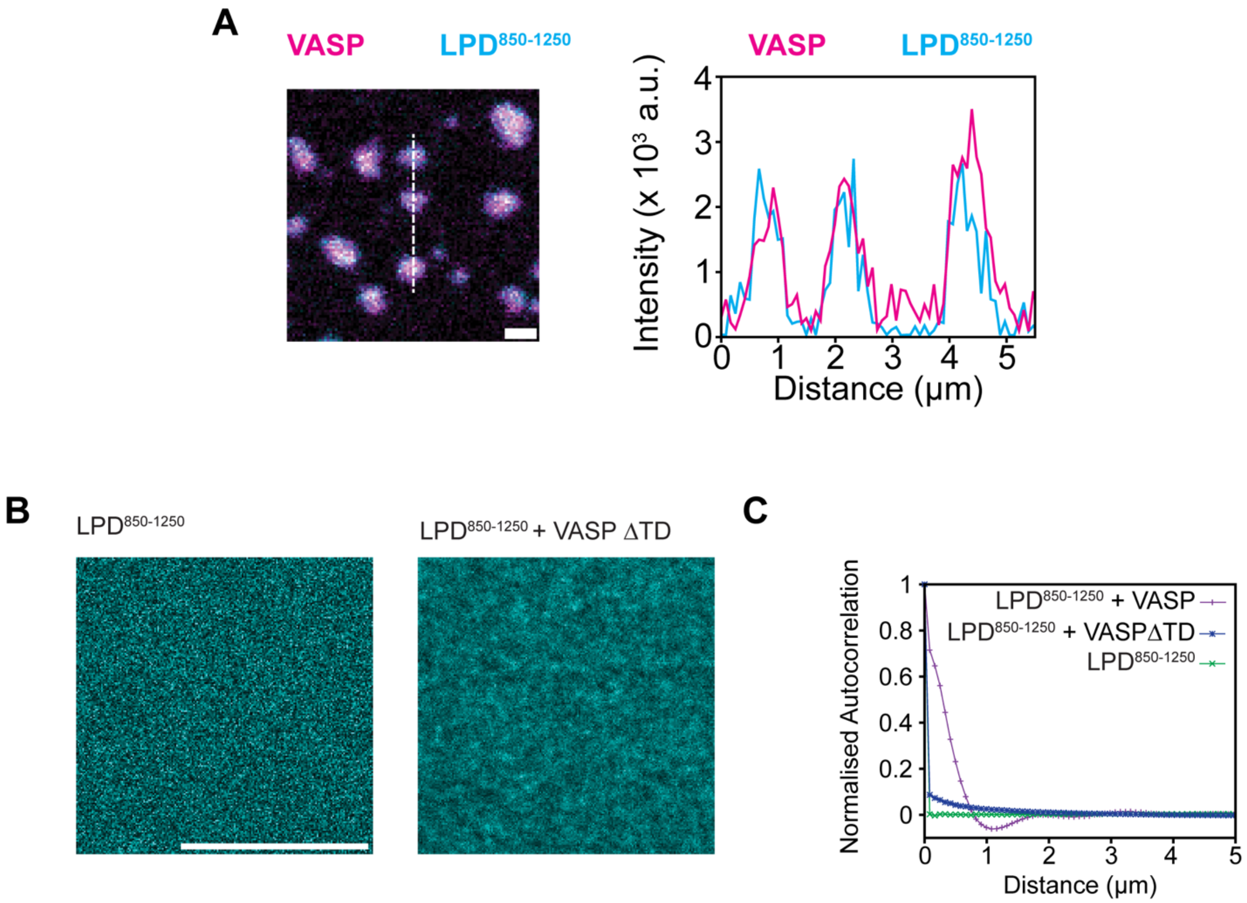


**Supplementary Figure 3: Phase separation and clustering of LPD$^{850\text{-}1250}$ with VASP and VASP-ΔTD on SLBs.**

**A**) Line intensity profile (**Right**) along dashed line (**Left**) drawn across LPD$^{850\text{-}1250}$ -VASP clusters (100 nM LPD$^{850\text{-}1250}$ and 2.5 μM VASP) on a supported lipid bilayer.
**B**) Representative image of 100 nM His tagged LPD$^{850\text{-}1250}$ coupled supported lipid bilayer (1% Ni-NTA, 99 DOPC) before (**Left**) and after (**Right**) incubation with 2.5 μM VASP-ΔTD (unlabeled). Scale bar 5 μm
**C**) Auto-correlation curves of LPD$^{850\text{-}1250}$, by itself, with VASP and VASP-ΔTD corresponding to concentrations mentioned in **A** and **B**.

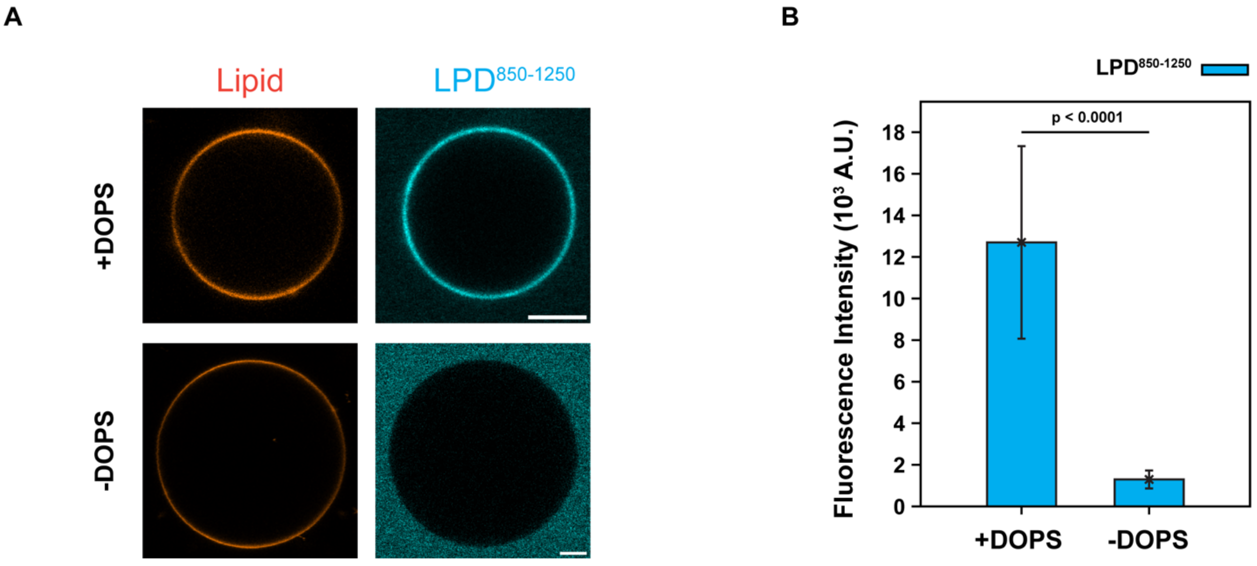


**Supplementary Figure 4: DOPS dependent recruitment of LPD$^{850\text{-}1250}$ to GUVs**

**A:** Representative images of LPD$^{850\text{-}1250}$ (1 μM doped with 100 nM Alexa 647 labeled protein) recruited to giant unilamellar vesicles (with and without 45% DOPS). Scale Bar 5 μm.

**B:** Quantification of fluorescence intensity of LPD$^{850\text{-}1250}$ on GUVs corresponding to conditions shown in **A**. Error bar represents +/- 1 standard deviation. N > 10 GUVs. Statistical analysis 2 tailed unpaired student t test.

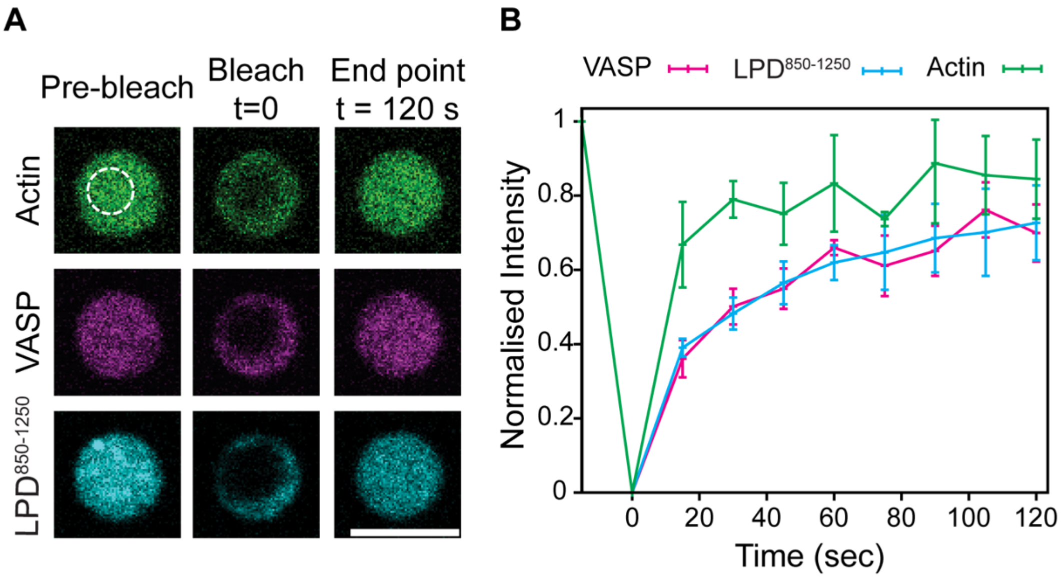


**Supplementary Figure 5: FRAP recovery of Actin VASP and LPD$^{850\text{-}1250}$ in condensates treated with LatA.**

**A** Image series and **B** normalised fluorescence recovery curves of LPD$^{850\text{-}1250}$-VASP condensate and actin (20 µM VASP, 10 µM LPD$^{850\text{-}1250}$ and 5 µM Actin) treated with LAT-A (+ 5µM LatA). Experiment performed 30 minutes after the introduction of LatA. The bleached area is enclosed by the dashed white circle. Scale bar 5 µm.

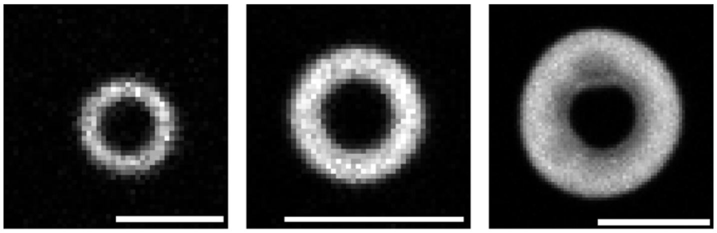


**Supplementary Figure 6: Instances of actin rings of various sizes**

Actin polymerized in LPD$^{850-1250}$-VASP condensates Representative images of actin (5 µM with 250 nM Rhodamine green actin) polymerized in LPD$^{850-1250}$-VASP condensates prepared by mixing 10 µM LPD$^{850-1250}$ with 20 µM VASP. Images corresponding to the low (2.67 µm), medium (3.8 µm) and high (7.22 µm) diameter of actin rings observed in the system after 30 minutes of polymerization are shown here.

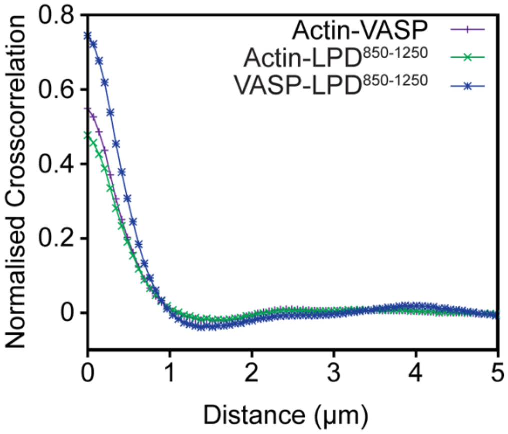

**Supplementary figure 7: Cross-correlation of LPD$^{850\text{-}1250}$, actin and VASP signal on SLBs**

Cross correlation plots of the pair-wise fluorescent channels (LPD$^{850\text{-}1250}$ -VASP, VASP-actin and actin- LPD$^{850\text{-}1250}$) corresponding to Actin (1 µM actin with 250 nM Rhodamine green Actin) polymerized on a supported lipid bilayer with preformed His tagged LPD$^{850\text{-}1250}$ -VASP clusters (100 nM Alexa 647 LPD$^{850\text{-}1250}$, 2.5 µM VASP with 250 nM Alexa 594 VASP respectively).

**A) Replotting data from 4B**

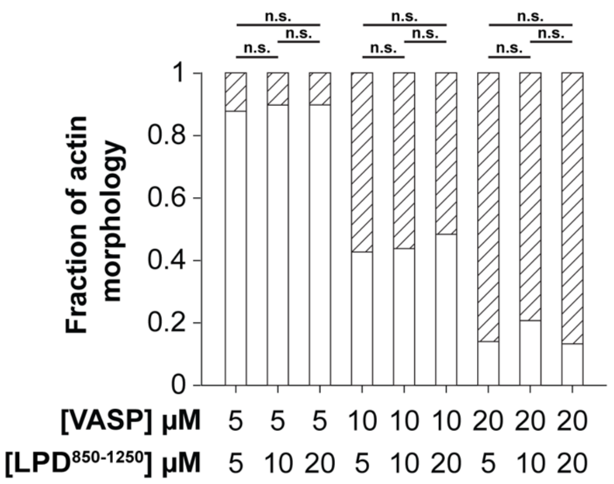


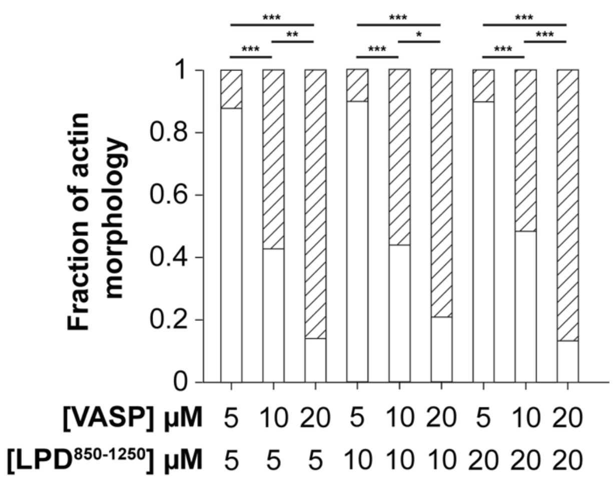


**B) Replotting data from 4D**

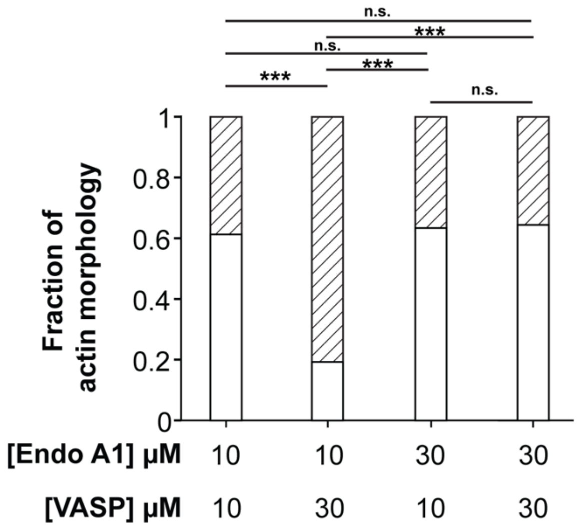


**Supplementary figure 8: Statistical analysis of actin morphologies obtained from Lamellipodin$^{850\text{-}1250}$-VASP and Endophilin-VASP condensates.**

**A)** Data shown in Fig **4B**, replotted as is with VASP concentration kept constant (Left), replotted with constant Lamellipodin concentration (Right).

**B)** Data shown in Fig **4D** replotted as is.

Pairwise Chi squared test performed, *, **, *** correspond to Bonferroni adjusted p value of <0.05, <0.001 and <0.0001. n.s means not significant.

A

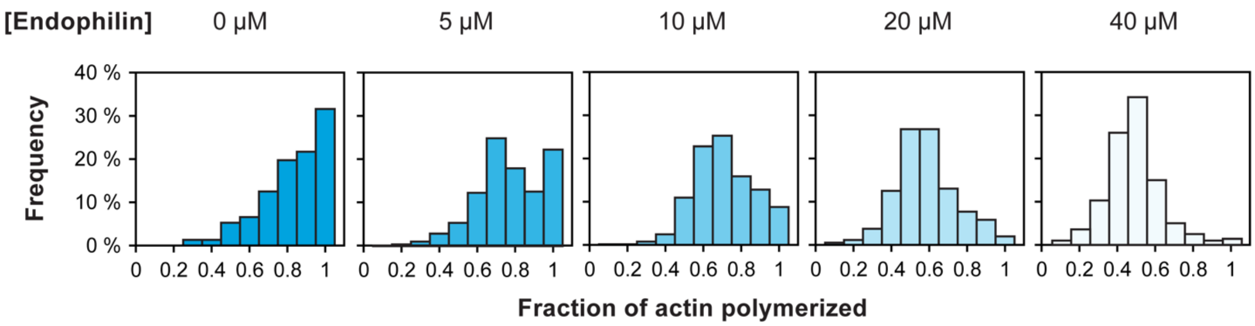


B

| Endophilin | VASP | LPD$^{850-1250}$ | Average actin polymerized | Average actin polymerized ratio w.r.t to 0 µM Endophilin | Relative loss of actin polymerization efficacy % |
|---|---|---|---|---|---|
| 0 µM | 10 µM | 10 µM | 0.78 | 1 | 0 |
| 5 µM | 10 µM | 10 µM | 0.72 | 0.92 | 8 |
| 10 µM | 10 µM | 10 µM | 0.66 | 0.84 | 16 |
| 20 µM | 10 µM | 10 µM | 0.53 | 0.68 | 32 |
| 40 µM | 10 µM | 10 µM | 0.43 | 0.55 | 45 |

Example Calculation, using 10 µM, 10 µM, 10 µM Endophilin, VASP, LPD$^{850-1250}$ polymerization data

Average actin poltmerized = 0.66

Ratio w.r.t 0 µM Endophilin = 0.66 / 0.78 = 0.84

Relative loss in actin polymerization efficacy = 1 - 0.84 = 0.16 or 16 %

C

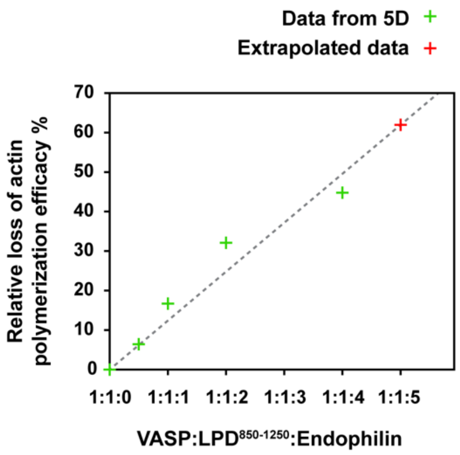


Fit line Equation y = 12.4 x,

Relative loss of actin polymerization efficacy(Endophilin ratio) = 12.4 *(Endophilin ratio)

Expected loss of actin polymerization at 1:1:5 VASP:Lamellipodin$^{850-1250}$:Endophilin = 62%

**Supplementary figure 9: Quantification of the effect of Endophilin on the extent of actin polymerization in LPD$^{850\text{-}1250}$-VASP condensates**

**A** Histogram depicting the distribution of actin polymerization efficiencies in LPD$^{850\text{-}1250}$-VASP-EndoA1 (10 µM, 10 µM, 0-40 µM) corresponding to **Figure 5D Left** in main text.

**B** Table showing the data and an example (below) depicting the process used to calculate the relative extent of loss of actin polymerization for data plotted in **A**.

**C Top** Plot showing the relationship between the relative loss of actin polymerization efficacy and the ratio of EndoA1 w.r.t LPD$^{850\text{-}1250}$ and VASP in the condensate. Dashed line corresponds to the best fit. **Bottom** Extrapolation of data to estimate the loss of actin polymerization for 1:1:5 LPD$^{850\text{-}1250}$-VASP-EndoA1.

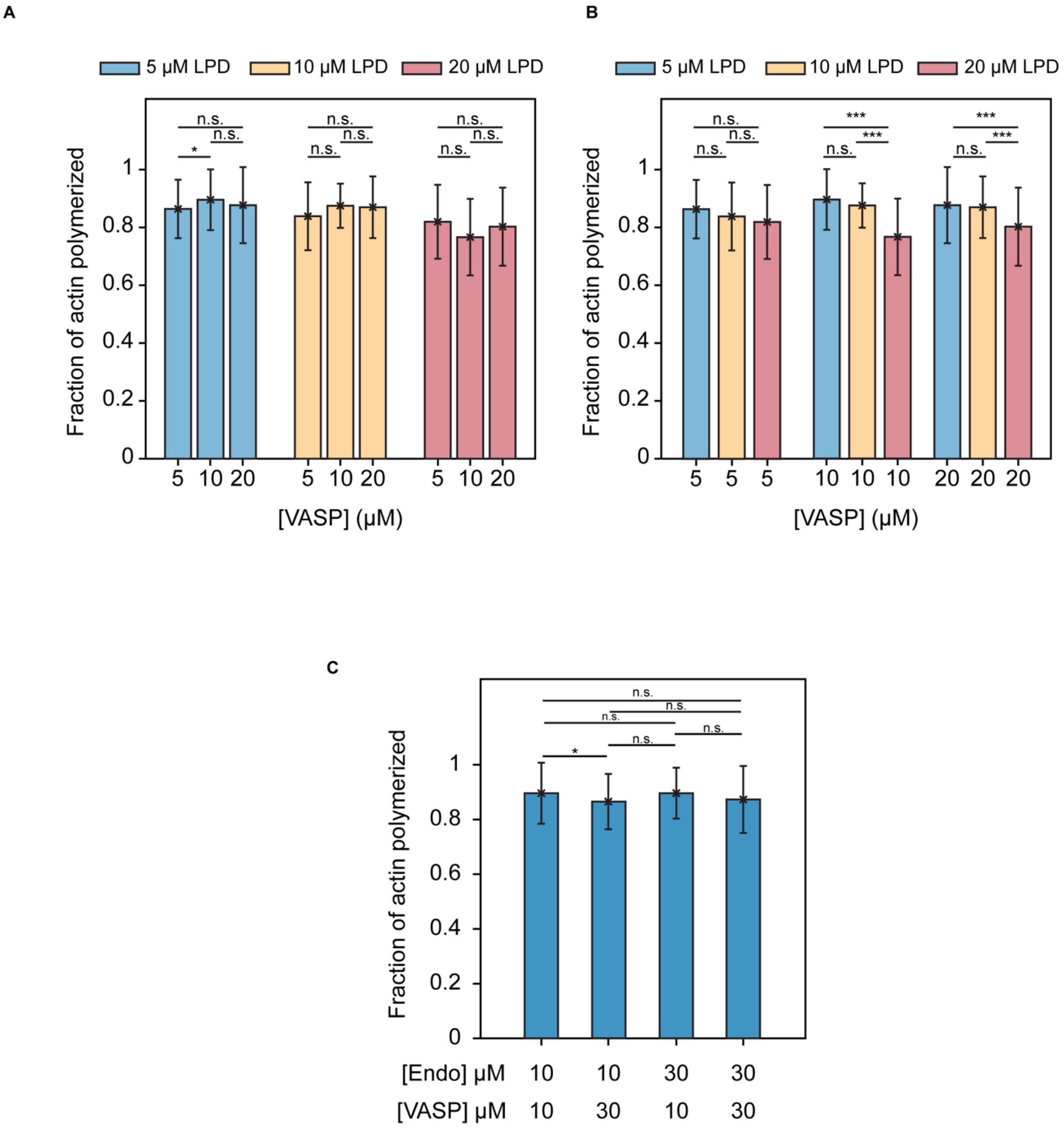


**Supplementary figure 10:** Quantification of degree of polymerization of actin in A,B) Lamellipodin$^{850-1250}$-VASP (grouped by constant Lamellipodin$^{850-1250}$ (A) and VASP (B) concentration), and C) Endophilin-VASP condensates determined by calculating the overlap between Phalloidin-A647 and Rhodamine green actin. Mean +/- S.D. shown in bar graph. For A,B) Left to right n = 56, 88, 43, 55, 38, 47, 47, 59, 304. For C) Left to right n = 151, 166, 46, 40. Pairwise Chi squared test performed, *, *** correspond to Bonferroni adjusted p value of <0.05 and <0.0001. n.s means not significant.

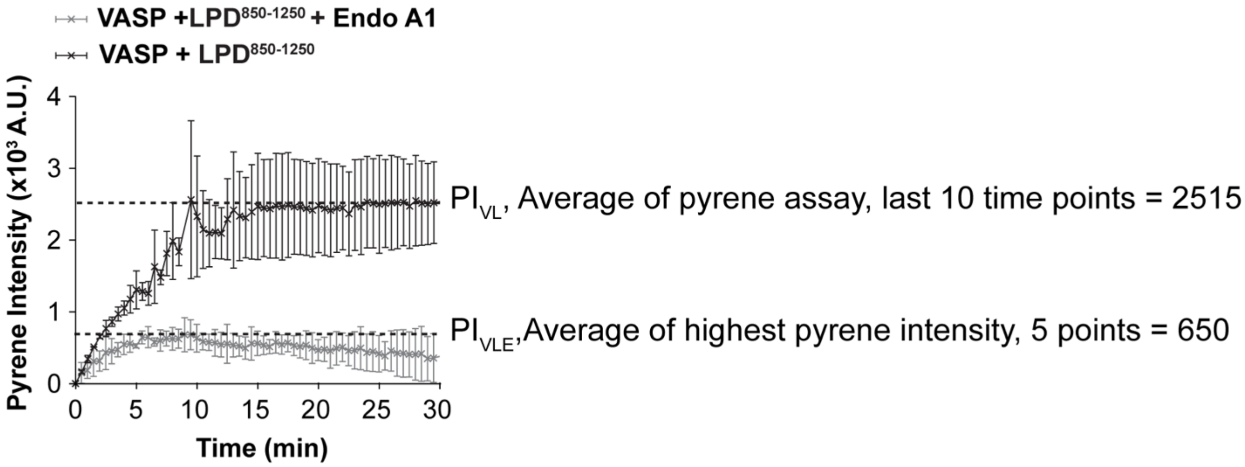


$$\text{Relative actin polymerization} = \frac{PI_{VLE}}{PI_{VL}} = \frac{650}{2515} = 0.25$$

Loss of relative actin polymerization = 1 - 0.25 = 0.75 or 75 %

**Supplementary figure 11: Calculation of the loss of actin polymerization using pyrene fluorescence data shown in Fig 6A.**
Average fluorescence signal obtained for the last 10 time points for VASP+ LPD$^{850\text{-}1250}$ and 5 time points for VASP+ LPD$^{850\text{-}1250}$+Endophilin used calculating ratio of actin polymerized.

PPPSPLSPVPSVVKQIASQFPPPPTPPAMESQPLKPVPANVAPQSPPAVKAKPKWQP
SSIPVPSPDFPPPPPPESSLVFPPPPPPSPVPAPPPPPPPTASPTPDKSGSPGKKTSKTSS
PGGKKPPPTPQRNSSIKSSSGAEHPEPKRPSVDSLVSKFTPPAESGSPSKETLPPPAA
PPKPGKLNLSGVNLPGVLQQGCVSAKAPVLSGRGKDSVVEFPSPPSDSDFPPPPPPET
ELPLPPIEIPAVFSGNTSPKVAVVNPQPQQWSKMSVKKAPPPTRPKRNDSTRLTQAEIS
EQPTMATVVPQVPTSPKSSLSVQPGFLADLNRTLQRKSITRHGSLSSRMSRAEPTATM
DDMALPPPPPPELLSDQQKAGYGGSHISGYATLRRGPPPAPPKRDQNTKLSRDW

**Supplementary figure 12**: **Sequence of Lamellipodin$^{850\text{-}1250}$** with Endophilin binding sites highlighted in red and VASP binding sites highlighted in yellow. Positively charged residues shown in orange.

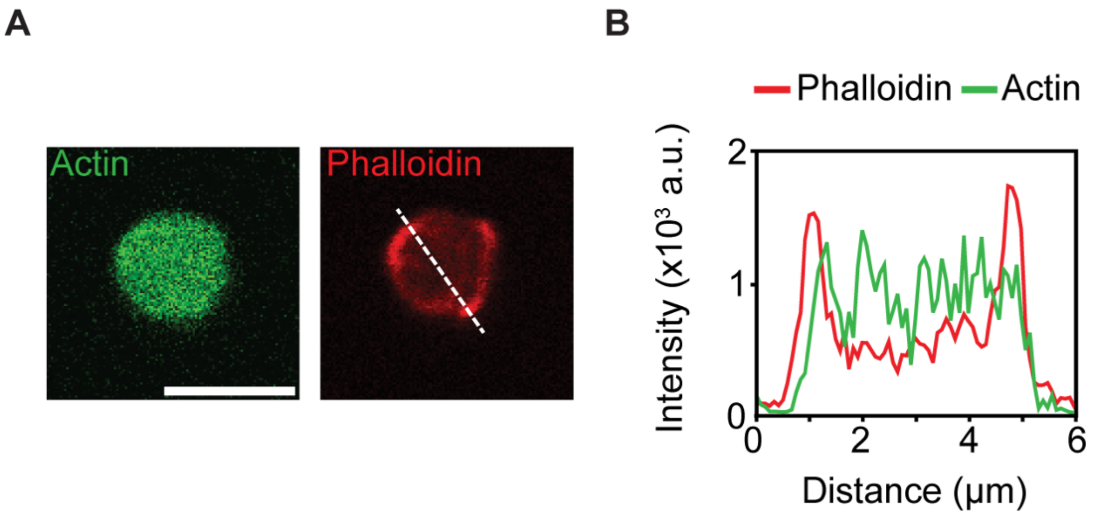


**Supplementary figure 13: Phalloidin staining of actin polymerized in LPD$^{850-1250}$-VASP-Endophilin condensates.**

**A** Staining filamentous actin polymerized in Lamellipodin$^{850-1250}$-VASP-Endophilin condensates (10μM, 20 μM, 40 μM respectively) using phalloidin-A647. Images acquired after 30 minutes of phalloidin treatment. Scale bar 5 μm. **B** Line intensity profile along white dashed line shown in image.

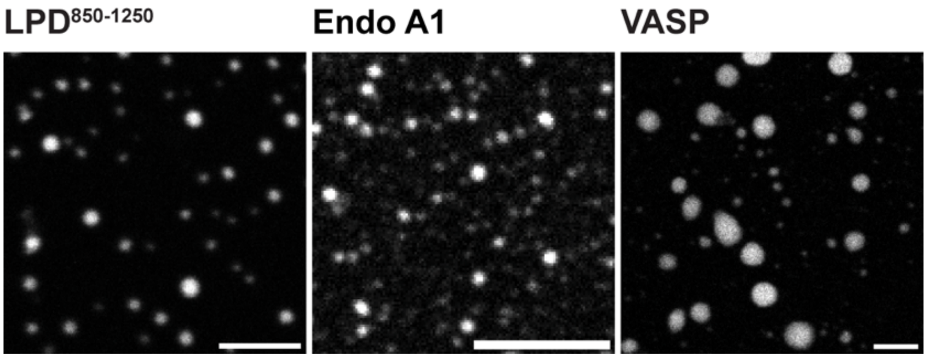


**Supplementary figure 14: Protein condensates formed by LPD$^{850-1250}$, Endophilin or VASP.**
Representative images depicting the formation of single protein condensates of (Left to Right) Lamellipodin$^{850-1250}$, Endophilin and VASP (20 µM total 1.25% labeled protein) formed in the presence of 5% PEG. Scale bar 5 µm.
